# SlCIPK26 enhances tomato fertility by activating the K^+^ transporter SlHAK5 in reproductive tissues

**DOI:** 10.64898/2026.05.12.724518

**Authors:** Almudena Martínez-Martínez, Adela Belchí, Elisa Jiménez-Estévez, Alberto Lara, Adrián Yáñez, Vicente Martínez, Francisco Rubio, Manuel Nieves-Cordones

## Abstract

In tomato plants, the potassium (K⁺) transporter SlHAK5 is integral to root K⁺ uptake and overall plant fertility. Under K⁺ deficiency, *SlHAK5* expression is induced in roots and the encoded transporter is activated via the Ca²⁺-sensing CIPK/CBL complex SlCIPK23/SlCBL1-9. In Arabidopsis, multiple CIPK/CBL complexes can activate AtHAK5, providing alternative regulatory pathways that enhance K⁺ uptake. However, the architecture of CIPK/CBL signaling networks has diverged among plant species, necessitating species-specific identification of novel regulatory components. Accordingly, we screened additional tomato CIPK proteins for their capacity to modulate SlHAK5 activity in yeast. SlCIPK15 and SlCIPK26 emerged as potent activators of SlHAK5, acting in concert with SlCBL9. Functional characterization of *slcipk15* and *slcipk26* mutants revealed that neither contributed significantly to SlHAK5-mediated K⁺ uptake in roots. Conversely, both mutants exhibited impaired pollen tube elongation, correlating with reduced K⁺ content in pollen relative to wild type. Notably, *slcipk26* mutants displayed more severe pollen defects, phenocopying the *slhak5* mutant. Further analyses demonstrated that *slcipk26* plants suffered compromised seed set and pistil function, paralleling the reproductive deficiencies observed in *slhak5* mutants. These findings implicate SlCIPK26 as the principal regulator of SlHAK5 in reproductive tissues. Collectively, our data underscore the role of CIPK paralogs in orchestrating tissue-specific regulation of target proteins, thereby enabling fine-tuned modulation of K⁺ transport essential for both vegetative and reproductive development.

## Introduction

Over the course of evolution, plants have established sophisticated molecular, cellular, and physiological systems that allow them to perceive stress signals, transmit this information internally, and initiate appropriate responses. These responses include adjustments in gene expression, metabolic pathways, and developmental processes, which collectively enhance their resilience to stresses such as drought, salinity, and nutrient deficiencies (Lamers et al., 2020). A key component of this adaptive response is the modulation of ion transport systems by calcineurin B-like protein (CBL)–interacting protein kinase (CIPK) networks, a conserved regulatory mechanism across land plants that confers versatility and specificity to signal transduction pathways (Tang et al., 2020a).

Potassium (K^+^) is an essential macronutrient for plants, and its uptake from the soil solution, followed by its distribution throughout the plant, constitutes a fundamental physiological process (Coskun and White, 2023). This process necessitates the transmembrane transport of K^+^ mediated by specialized transporters and is tightly regulated through complex signaling networks (Nieves-Cordones and Rubio, 2025). In tomato (*Solanum lycopersicum*), the SlHAK5 transporter is pivotal for K^+^ nutrition, particularly under conditions of low external K^+^ availability. Evidence for its critical role is provided by the impaired growth observed in *slhak5* mutant plants subjected to K^+^-deficient conditions (Nieves-Cordones et al., 2020). Beyond root uptake, SlHAK5 contributes to tomato fertility. In *slhak5* plants, defects in both pollen and pistil tissues resulted in the formation of parthenocarpic (seedless) fruits due to impaired fertilization (Jesus Amo et al., 2024; Nieves-Cordones et al., 2020).

Regulation of SlHAK5 activity occurs at both transcriptional and post-translational levels. In roots, the expression of *SlHAK5* is strongly upregulated under K^+^ starvation, while its transporter activity is modulated post-translationally by the SlCIPK23-SlCBL1/9 complexes (Amo et al., 2021; Nieves-Cordones et al., 2008, 2007). CIPK-CBL complexes form elaborate interaction networks with target proteins, enabling a single transport system to be regulated by multiple CIPK-CBL combinations. For example, in *Arabidopsis thaliana*, AtCIPK23, AtCIPK1, and AtCIPK9 cooperatively activate AtHAK5 in conjunction with CBL1 (Lara et al., 2020), while the K^+^ channel AKT1 is activated by AtCIPK23, AtCIPK6, and AtCIPK16 (Sung et al., 2007). Notably, individual CIPKs can regulate multiple ion transport systems; for instance, SlCIPK23 activates both the SlHAK5 and LKT1 K^+^ transport systems while inhibiting the SlSKOR K^+^ channel in tomato plants (Amo et al., 2021; Nieves-Cordones et al., 2023). Beyond ion transport regulation, CIPKs also participate in broader physiological processes. Arabidopsis CIPK23, for example, interacts with TAP46—a direct target of the developmental regulator Target of Rapamycin (TOR) (Deng et al., 2020)—and with the blue light receptor kinases Phot1 and Phot2 (Inoue et al., 2020).

Regarding SlHAK5 regulation in tomato, aside from the established SlCIPK23/SlCBL1-9 and SlCIPK9/SlCBL1-9 complexes, no additional activating complexes had been identified prior to this study. The present investigation aimed to uncover novel SlCIPK kinases and Ca^2+^ sensors (SlCBLs) that modulate SlHAK5 function. The results demonstrate that the SlCIPK15/SlCBL1-9 and SlCIPK26/SlCBL9 complexes act as positive regulators of SlHAK5 in yeast. Functional characterization *in planta* revealed that both SlCIPK15 and SlCIPK26 contribute to the regulation of K^+^ uptake in pollen but not in roots. Significantly, *slcipk26* mutant plants exhibit reproductive tissue defects analogous to those observed in *slhak5* mutants, implicating SlCIPK26 as the principal regulator of SlHAK5 in reproductive organs.

## Materials and methods

### Plant material and growth conditions

Tomato plants (*Solanum lycopersicum* L. var. Micro-Tom) were used for physiological experiments. Seeds were sown in 1/5 Hoagland solid plates (0.8% agar). When seeds germinated, they were transferred to containers with a modified 1/5 Hoagland solution with the following macronutrients (mM): 1.4 Ca(NO_3_)_2_, 0.35 MgSO_4_, and 0.1 Ca(H_2_PO_4_)_2_ and micronutrients (μM): 50 CaCl_2_,12.5 H_3_BO_3_, 1 MnSO_4_, 1 ZnSO_4_, 0.5 CuSO_4_, 0.1 H_2_MoO_4_, 0.1 NiSO_4_, and 10 Fe-EDDHA. KCl (1.4 mM) was added to the +K solutions, whereas no KCl was added to the –K solutions. Plants were grown in the –K solution for 7 d. For pollen and fructification experiments, tomato plants were grown in pots containing peat (Composana, Compo Expert) for 4-8 weeks. Tomato plants were grown in a controlled environment growth chamber with a 16/8 h light/dark photoperiod. The plants were grown at 25 °C (day) and 20 °C (night) temperatures, 65% relative humidity, and 360 μmol m^−2^ s^−1^ flux density. The pH of the nutrient solution was adjusted to 5.5 daily. The growth solution was changed weekly.

### Yeast growth and Rb^+^ uptake measurements

The *trk1 trk2* 9.3 yeast strain (*MATa, ena1Δ::HIS3::ena4Δ, leu2, ura3, trp1, ade2, trk1Δ, trk2::pCK64*), deficient in its endogenous K^+^ uptake systems, TRK1 and TRK2, was used in this study. The cDNA encoding the previously described chimeric qSlHAK5 (Nieves-Cordones et al., 2008) was cloned into the pYPGE15 vector (Brunelli and Pall, 1993). The cDNAs encoding the tomato SlCBL1 and SlCBL9 were cloned in the p425GPD (Mumberg et al., 1995). In previous phylogenetic analyses, 29 *CIPK* genes were identified in the tomato genome (Edel and Kudla, 2015; Nieves-Cordones et al., 2023). cDNAs encoding these proteins, except for SlCIPK23 and SlCIPK9, were synthesized and cloned into the yeast pESC TRP vector (GenScript, Piscataway, NJ, USA). The cDNA sequences used for gene synthesis are listed in Table S1. Plasmids were transformed into yeast cells as previously described (Elble, 1992). Synthetic SD medium (Sherman, 1991) was used for the selection of transformants. Minimal AP medium (Rodriguez-Navarro and Ramos, 1984) supplemented with either 50 mM or 0.1 mM KCl was used for yeast complementation assays. Ten microliter drops of serial dilutions of the yeast suspensions were deposited on plates with solid AP medium with the indicated K^+^ concentrations and incubated at 28 °C 2-5 d.

For Rb^+^ uptake experiments, yeast expressing the constructs indicated in each figure were grown overnight in AP medium supplemented with 5 mM K^+^. The cells were collected by centrifugation and washed with distilled water. They were then grown in AP medium with no added K^+^ for 4h. After a new centrifugation step, the cells were incubated in AP medium supplemented with 0.1 mM RbCl. Samples were obtained at different time intervals, centrifuged, washed with cold double-distilled water, and suspended in 0.1 M HCl. Rb^+^ was measured from the supernatant solution using a Perkin Elmer AAnalyst 400 spectrometer. Rb^+^ uptake rates were determined from the internal Rb^+^ accumulated in yeast cells per dry weight and time.

### K^+^ and Rb^+^ content measurements in plants

At the end of the growth period, +K and –K plants were transferred to a solution containing 20 µM RbCl for 6 h. The plants were then separated into roots and shoots, and their fresh and dry weights were determined. The dried plant material was digested with HNO_3_:H_2_O_2_ (5:3 v:v) in a microwave (CERM MarsXpress, North Carolina, USA), and the K^+^ content was determined by inductively coupled plasma (ICP) mass spectrometry using an Iris Intrepid II ICP spectrometer (Thermo Electron Corp., Franklin, MA, USA). Rb^+^ uptake rates were calculated based on the accumulation of Rb^+^ in the plant tissues per unit of time and per unit of root dry weight.

### Real time qPCR

Plant tissues were frozen in liquid nitrogen and then RNA was extracted using the Macherey-Nagel NucleoSpin RNA Plant Kit (Neumann-Neander, Germany). DNA-free kit (Thermo Fisher Scientific, Waltham, MA, USA) was applied to the RNA samples to eliminate contaminating DNA. A 7500 Real-Time PCR System (Thermo Fisher Scientific) was used for Real-time PCR using the default cycle settings. The expression levels of *SlCIPK15* and *SlCIPK26* were calculated by taking into account the expression level of the housekeeping gene encoding the elongation factor SlEF1α and expressed as 40 − ΔCt (ΔCt = Ct *target* − Ct *SlEF1α*) (Nieves-Cordones et al., 2020). The primer sequences are listed in Table S2.

### Gene edition with CRISPR-Cas

To edit the *SlCIPK15* (Solyc05g052270) and *SlCIPK26* (Solyc11g062410) genes, two sgRNAs were designed for each gene (Figure S1A and S1B) using the Breaking-Cas tool (Oliveros et al., 2016). As *SlCIPK15* has only one exon, both sgRNAs targeted this exon (Figure S1A). In the case of *SlCIPK26*, sgRNAs targeted exons 1 and 7 (Figure S1B). Target sequences were selected because of their high specificity (cut probability in off-target sites <0.1%). Cloning protocols and plasmids were based on Golden Braid technology (Vazquez-Vilar et al., 2021). The two sgRNAs (Table S2) were assembled to produce polycistronic RNA (GB1207 and GB1208 parts) in the plant. The plant expression vector pDGBΩ3 contained a kanamycin resistance gene, Ds-RED marker expressed in seeds, Cas9 nuclease (GB2234 part), and the sgRNA transcriptional unit. The resulting construct was introduced into *Agrobacterium tumefaciens* strain LBA4404. Plant transformation was carried out by the Plant Transformation Service of CEBAS-CSIC by infecting the cotyledons of WT Micro-Tom with *Agrobacterium* suspensions and subsequent in vitro culture procedures. Genomic DNA flanking the target regions of the sgRNA was amplified by PCR and sequenced. Subsequently, the allele sequences were deduced using the DECODR tool (Bloh et al., 2021). Plant lines harboring frameshift mutations in target genes were selected for further characterization. Two independent lines were obtained for each mutant (Figure S1C and S1D, respectively). Plant genotypes were determined in T1 and T2 plants, and all experiments were carried out with plants from these generations.

### Determination of germination, tube length and K^+^ content of pollen

When tomato plants were four weeks old, pollen was harvested from flowers onto microscope slides containing germination medium (0.5 % agar, 10% sucrose, and 50 ppm H_3_BO_3_). The K^+^ concentration in the germination medium was 100 µM K^+^. Pollen was incubated for 4 h inside a Petri dish at 25°C and then observed under a Leica DM6 microscope. If the pollen tube was >5 μm in length, it was labelled as having germinated. The internal free K^+^ concentration of pollen grains was estimated using the Ion-K^+^ green 4 a.m. dye (IPG-4, Ion Biosicences, San Marcos, TX USA) at a concentration of 20 μM in the germination medium. The pollen was incubated for 16 h at 25 °C in the presence of the dye. Samples were observed under a Leica DM6 epifluorescence microscope, and the fluorescence signal was recorded with a 527/30 nm filter after excitation at 480/40 nm. ImageJ software was used to determine the pollen tube length and IPG-4 fluorescence. The mean fluorescence of pollen grains without dye (pollen autofluorescence) was subtracted from the fluorescence signal in the IPG-4-incubated samples.

### BiFC assays in Nicotiana benthamiana

The cDNAs encoding *SlCIPK15*, *SlCIPK26*, and *AtSOS1* were introduced into pSPYNE173(R), whereas the cDNA encoding *SlHAK5* was cloned into pSPYCE(MR) (primers are listed in Table S2). *A. tumefaciens* GV3101 strain was transformed with the indicated constructs and used for co-infiltration experiments on *Nicotiana benthamiana* leaves. *A. tumefaciens* strains were infiltrated at an optical density of 0.25 at 600 nm wavelength. The P19 helper plasmid was added to all mixtures at an optical density of 0.1 at 600 nm. A LEICA DM6 microscope was used to observe the fluorescence with an L5 filter (excitation BP480/40, Dic. Mirr.505, emission BP527/30).

## Results

### SlCIPK15 and SlCIPK26 as positive regulators of the K⁺ transporter SlHAK5

To identify novel CIPK regulators of the K⁺ transporter SlHAK5, a comprehensive yeast-based screen was conducted employing all tomato CIPKs except SlCIPK23 and SlCIPK9, previously established activators of the K⁺ transporter (Amo et al., 2021; Martínez-Martínez et al., 2024). The yeast strain 9.3, deficient in endogenous TRK transport systems and expressing the chimeric transporter qSlHAK5 alongside either SlCBL1 or SlCBL9 plasma membrane CBLs, was transformed individually with plasmids encoding 27 tomato CIPKs. Growth complementation assays were performed on AP plates supplemented with 0.1 mM K⁺. Positive controls included yeast expressing qSlHAK5+SlCIPK23+SlCBL1 or qSlHAK5+SlCIPK23+SlCBL9, while yeast strain 9.3 and qSlHAK5-expressing strains served as negative controls. All strains were also plated on AP medium containing 50 mM K⁺ to confirm baseline growth viability.

Following incubation, differential growth complementation was observed among strains on 0.1 mM K⁺ medium, ranging from no improvement relative to negative controls to levels comparable with positive controls (Figure 1). In yeast expressing SlCBL1, most strains exhibited poor growth at 0.1 mM K⁺, except for qSlHAK5+SlCIPK15+SlCBL1, which demonstrated robust complementation (Figure 1A). Although qSlHAK5+SlCIPK2+SlCBL1 showed enhanced growth relative to negative controls, it did not match the complementation conferred by SlCIPK15 (Figure 1A). In yeast expressing SlCBL9, qSlHAK5+SlCIPK26+SlCBL9 and qSlHAK5+SlCIPK15+SlCBL9 exhibited comparable growth levels, both approaching those of the positive control (Figure 1B). Constructs comprising qSlHAK5+SlCIPK2+SlCBL9, qSlHAK5+SlCIPK11+SlCBL9, and qSlHAK5+SlCIPK27+SlCBL9 yielded growth superior to negative controls, whereas other constructs resembled negative controls (Figure 1B).

**Figure 1.**
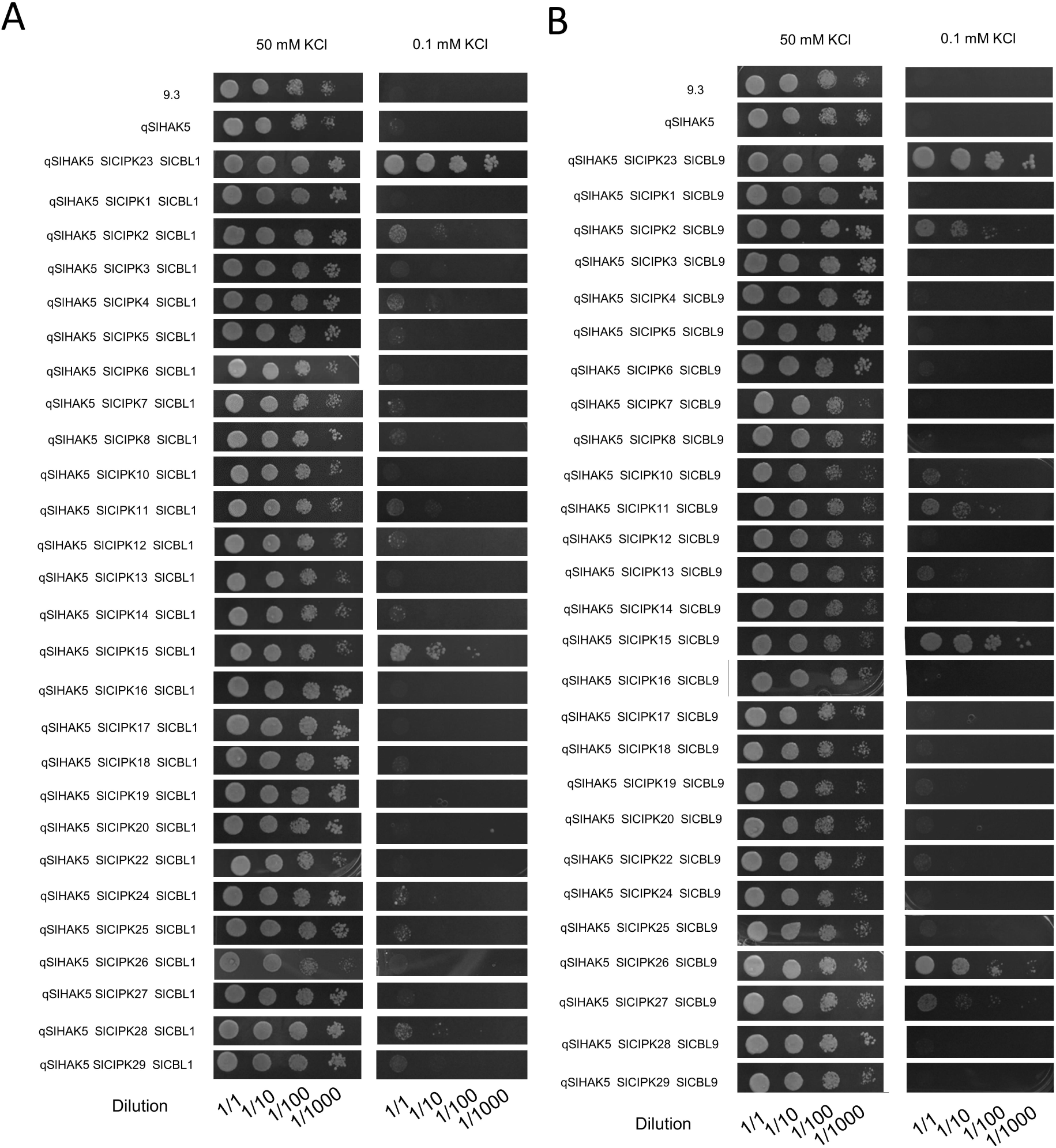
SlCIPK15 and SlCIPK26 activate qSlHAK5 in yeast. Growth tests for yeast expressing qSlHAK5, SlCBL1 or SlCBL9 and 28 tomato CIPKs. Vectors containing qSlHAK5, SlCBL1, or SlCBL9 and 28 tomato CIPKs were used to transform the 9.3 yeast strain, which cannot take up K^+^ from low external concentrations. Serial dilutions of yeast suspensions were incubated in AP minimal medium supplemented with 50 mM or 0.1 mM K^+^.

Integrating these findings from tomato with prior data from Arabidopsis (Lara et al., 2020), we identified CIPKs from both species that, in conjunction with CBL1/9, activate HAK5 transporters. Phylogenetic analysis was conducted to assess whether HAK5 activation correlated with protein sequence homology (Figure S2). This analysis revealed that certain homologous CIPKs from tomato and Arabidopsis exhibited conserved regulatory capacity on HAK5. Specifically, CIPK23, CIPK9, and CIPK26 were identified as activators. However, no consistent relationship between protein homology and HAK5 regulation was observed for other CIPKs; for instance, AtCIPK4 and AtCIPK7 enhanced AtHAK5 activity, whereas their tomato homologs SlCIPK4, SlCIPK7, and SlCIPK16 did not modulate SlHAK5 (Figure S2).

To validate that yeast growth complementation corresponded to qSlHAK5-mediated K⁺ uptake, short-term Rb⁺ uptake assays were conducted at an external concentration of 0.1 mM Rb⁺. Yeast expressing qSlHAK5+SlCIPK15+SlCBL9 and qSlHAK5+SlCIPK26+SlCBL9 exhibited rapid intracellular Rb⁺ accumulation (Figure 2A), with uptake rates of 2.76 ± 0.15 and 2.81 ± 0.22 nmol Rb⁺ mg DW⁻¹ min⁻¹, respectively (Figure 2B), representing 9- to 28-fold increases over the two negative controls. Nonetheless, SlCIPK15 and SlCIPK26 activated qSlHAK5 to a lesser extent than SlCIPK23, which mediated an uptake rate of 6.28 ± 0.54 nmol Rb⁺ mg DW⁻¹ min⁻¹ (Figure 2B). Additional assays of CIPKs yielding modest growth complementation (SlCIPK2, SlCIPK11, SlCIPK27) confirmed that qSlHAK5+SlCIPK2+SlCBL9 and qSlHAK5+SlCIPK11+SlCBL9 induced moderate Rb⁺ uptake increases (Figure 2C), albeit substantially below positive control levels (Figure 2D). qSlHAK5+SlCIPK27+SlCBL9 did not significantly differ from qSlHAK5 alone. Based on these results, SlCIPK15 and SlCIPK26 were selected for further characterization.

**Figure 2.**
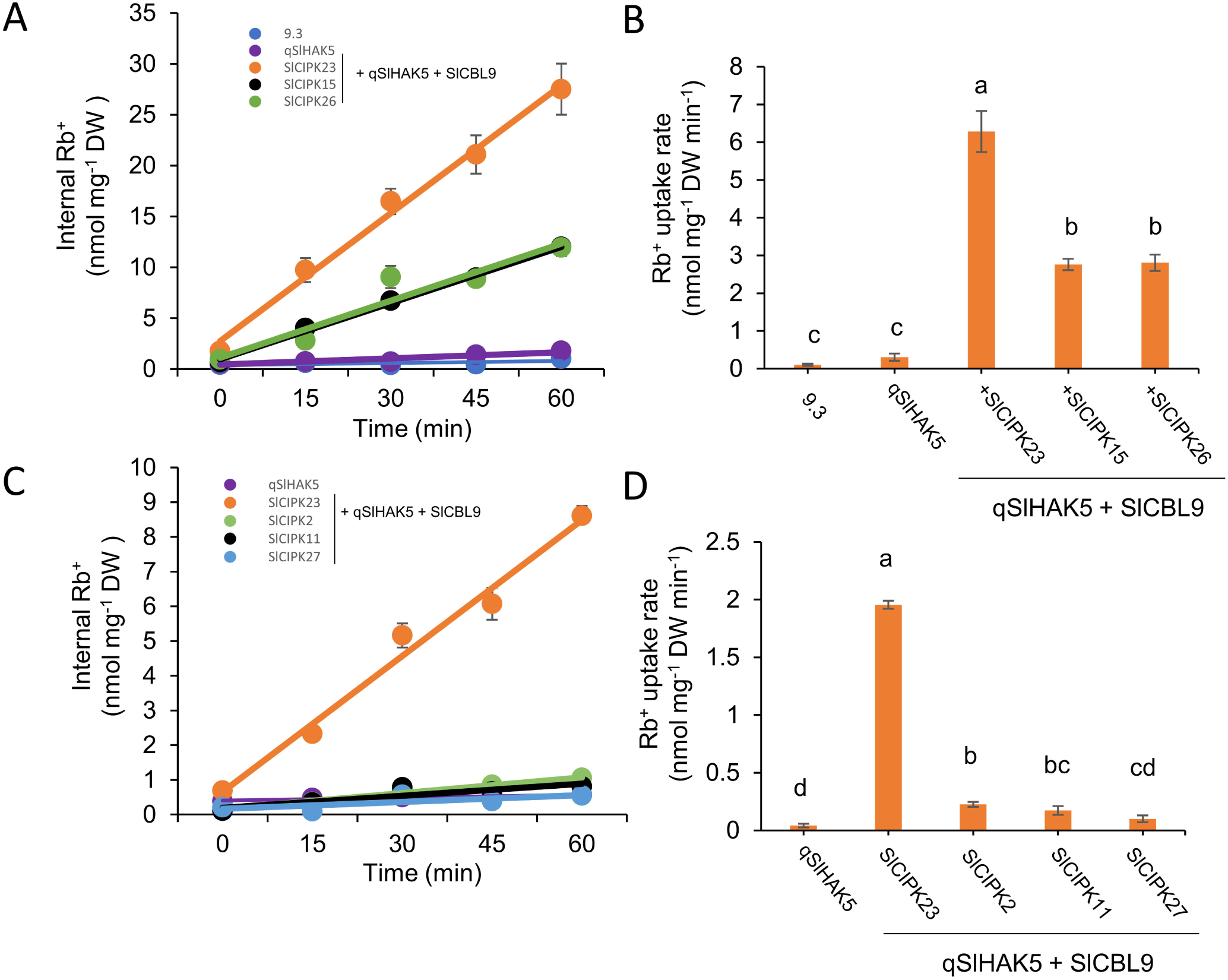
SlCIPK15 and SlCIPK26, together with SlCBL9, enhanced Rb^+^ uptake by qSlHAK5 in yeast. (a) Internal Rb^+^ content versus time from a 0.1 mM external Rb^+^ solution in K^+^-starved yeast cells expressing 9.3 transformed with empty vectors, qSlHAK5 alone, or in combination with SlCIPK23, SlCIPK15, SlCIPK26, and SlCBL9. (b) Rb^+^ uptake rates calculated after 60 min of the experiment, corresponding to the samples shown in (a). (c) Internal Rb^+^ content versus time from a 0.1 mM external Rb^+^ solution in K^+^-starved yeast cells expressing 9.3 transformed with empty vectors, qSlHAK5 alone, or in combination with SlCIPK23, SlCIPK2, SlCIPK11, SlCIPK27, and SlCBL9. (d) Rb^+^ uptake rates calculated after 60 min of the experiment corresponding to the samples shown in (c). Dots and bars show mean values of two repetitions, and error bars depict SE. Bars with different letters are significantly different according to the LSD test (p < 0.05).

To substantiate direct regulation of SlHAK5 by SlCIPK15 and SlCIPK26 via protein-protein interactions, bimolecular fluorescence complementation (BiFC) assays were performed in *Nicotiana benthamiana* leaves. SlHAK5 fused to the C-terminal half of YFP (YC) was co-expressed with AtSOS1-YN as a negative control (Figure 3A), and with SlCIPK15-YN or SlCIPK26-YN (Figures 3B, 3C). Three days post-Agrobacterium infiltration, YFP fluorescence was reconstituted in SlHAK5-SlCIPK15 and SlHAK5-SlCIPK26 samples, but not in the SlHAK5-AtSOS1 control, confirming specific interactions between SlHAK5 and these CIPKs.

**Figure 3.**
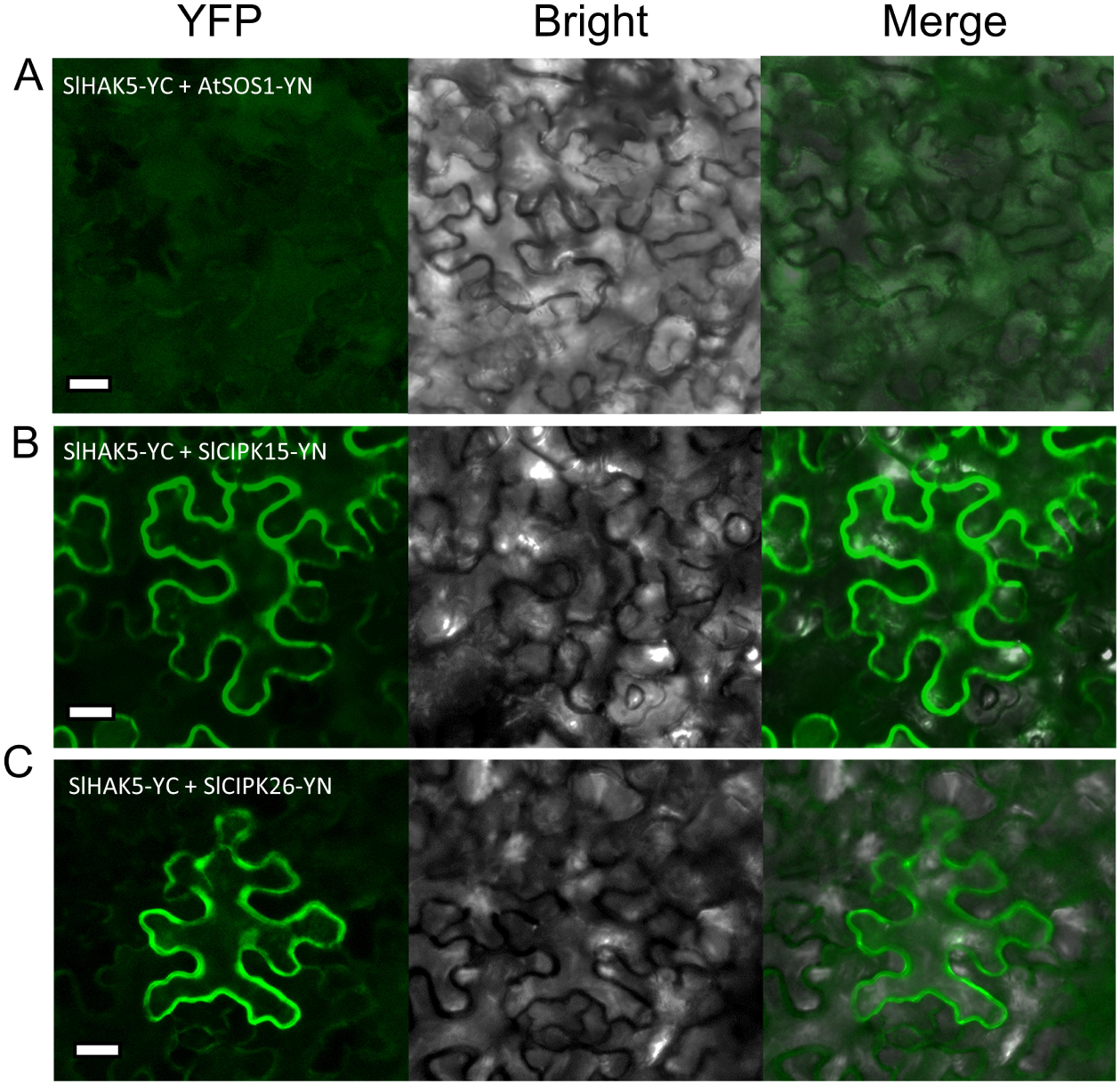
SlCIPK15 and SlCIPK26 interacted with SlHAK5 in BiFC assays. Each protein was fused to either the N-terminal (YFPN) or C-terminal (YFPC) half of the yellow fluorescent protein (YFP), and interactions were assessed upon co-expression of YFPN and YFPC protein fusions in *Nicotiana benthamiana* leaves. SlHAK5-YC and AtSOS1-YN were used as negative controls. For each construct, YFP signal, bright field, and merged images are shown. Scale bar = 25 μm.

### K⁺ uptake at low external concentrations is unaffected in *slcipk15* and *slcipk26* mutants

To elucidate the physiological roles of SlCIPK15 and SlCIPK26 *in planta*, CRISPR-Cas9-generated knockout mutants *slcipk15-1* and *slcipk26-1* were cultivated alongside wild-type (WT) plants under K⁺-sufficient and K⁺-deficient hydroponic conditions. K⁺ deficiency was imposed by withholding K⁺ from the nutrient solution for seven days prior to harvest. Rb⁺ uptake assays at 20 µM external Rb⁺—a concentration at which SlHAK5 accounts for 95% of root Rb⁺ uptake (Nieves-Cordones et al., 2020)—were conducted, alongside measurements of dry weight and tissue K⁺ content.

Under K⁺-sufficient conditions, root and shoot dry weights of *slcipk15-1* and *slcipk26-1* mutants were comparable to WT (Figure 4A). Under K⁺ deficiency, *slcipk15-1* exhibited marginally reduced shoot and root biomass, whereas *slcipk26-1* displayed reduced root dry weight relative to WT (Figure 4A). K⁺ content in *slcipk15-1* was similar to WT across all tissues and conditions (Figure 4B), while *slcipk26-1* showed elevated K⁺ in +K shoots and – K roots (Figure 4B). Rb⁺ uptake rates did not differ significantly between mutants and WT under either condition (Figure 4C). These findings were corroborated using independent mutant alleles (Figure S3A-C), indicating that SlCIPK15 and SlCIPK26 are dispensable for SlHAK5-mediated root K⁺ uptake.

**Figure 4.**
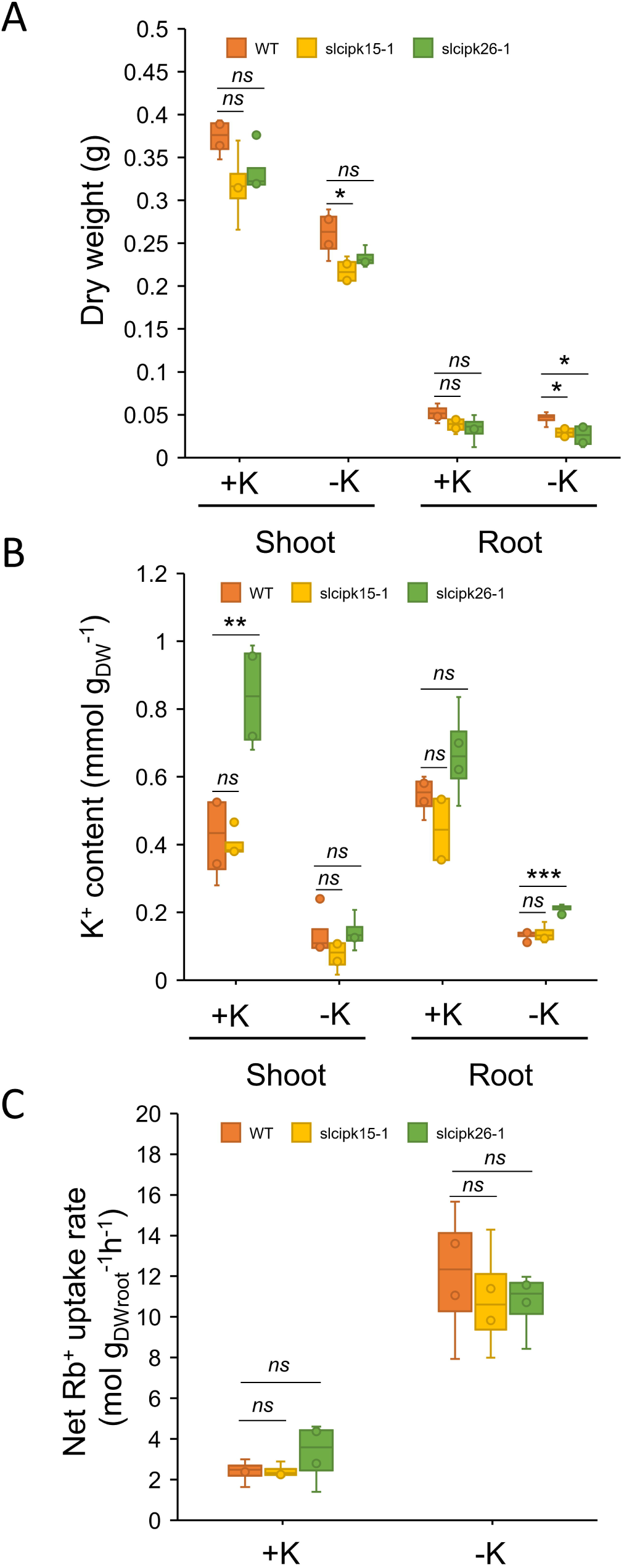
SlCIPK15 and SlCIPK26 did not contribute to Rb^+^ uptake at low external concentrations in tomato plants. Tomato plants were grown for 14d in +K solution, and a group of plants was starved of K^+^ for 7 d. On the last day of the experiment, a Rb^+^ uptake assay at 20 µM was performed, and the plants were harvested. (a) Dry weight, (b) K^+^ content, and (c) net Rb^+^ uptake rates of WT, *slcipk15-1*, and *slcipk26-1* plants grown under +K or –K conditions. Data are shown as boxplots (n = 4). *, ** and *** indicate p < 0.05, p < 0.01 and p < 0.001 in Student’s t test, respectively.

### The *slcipk26* mutant exhibits severe pollen defects

Despite the absence of root phenotypes, the possibility that SlCIPK15 and SlCIPK26 regulate SlHAK5 in other tissues was explored by quantifying their expression via qRT-PCR in vegetative and reproductive organs. *SlCIPK15* expression peaked in stamens, followed by petals, pistils, sepals, floral stems, leaves, stems, and roots (Figure 5A). *SlCIPK26* was most highly expressed in stamens, with substantial expression in leaves, petals, sepals, pistils, floral stems, stems, and roots (Figure 5B). These expression patterns were consistent with data from the Tomexpress database, which showed maximal expression of both CIPKs in flowers (Figure S4).

**Figure 5.**
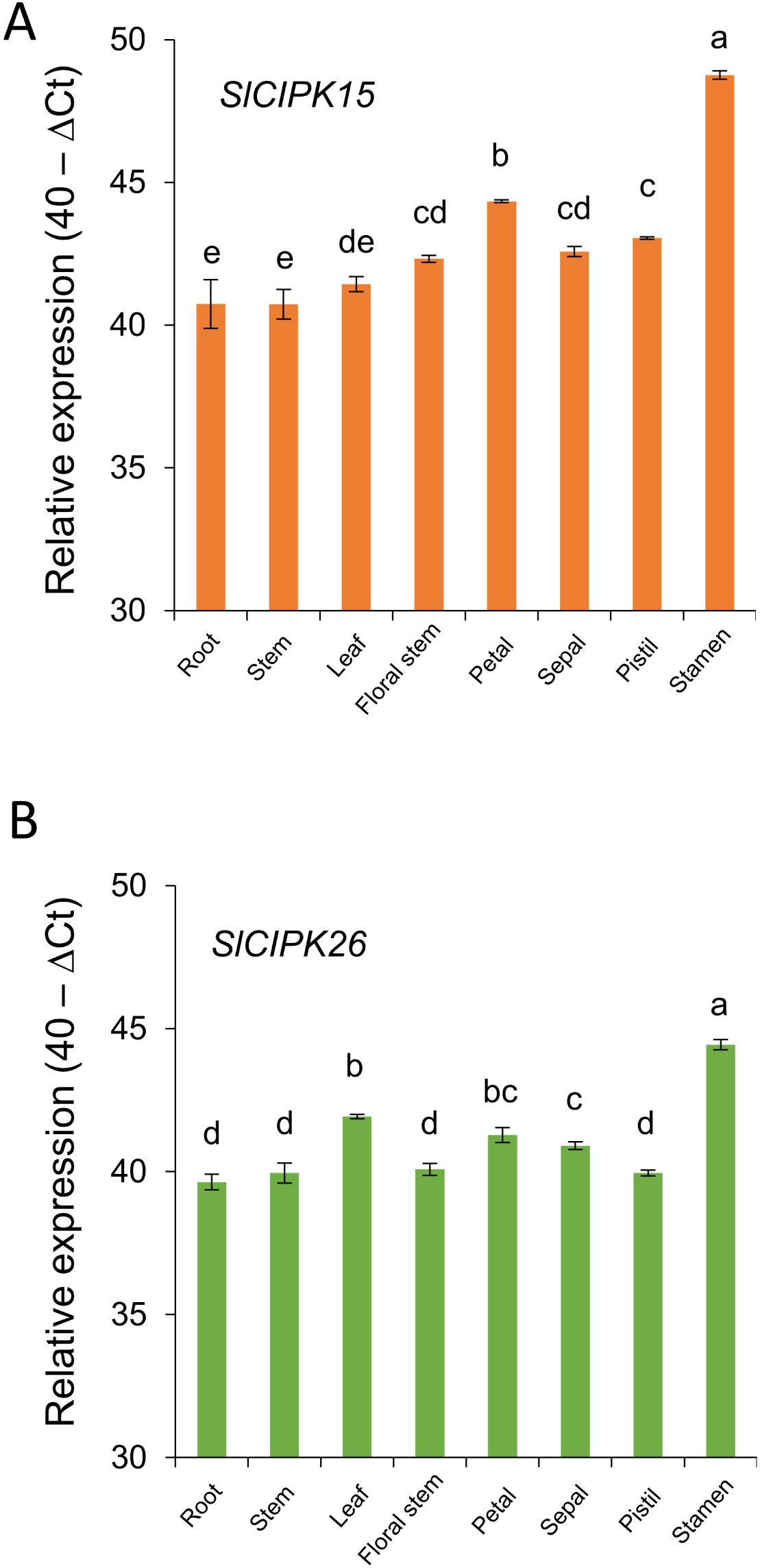
*SlCIPK15* and *SlCIPK26* were highly expressed in the stamens. WT plants were grown in +K solution and separated into different plant organs. Gene expression was quantified in cDNA samples by RT-qPCR using specific primers for *SlCIPK15* and *SlCIPK26*. Gene expression was calculated using the 40−ΔCt method. Data are mean ± SE (n = 2). Bars with different letters are significantly different at p < 0.05 according to the LSD test.

Given the elevated expression in stamens, *in vitro* pollen germination and tube elongation assays were conducted using pollen from WT, *slcipk15-1*, and *slcipk26-1* plants grown for four weeks. Two K⁺ concentrations were tested in germination media: 0.1 mM (no added K⁺) and 1 mM K⁺. *slcipk15-1* pollen exhibited normal morphology and germination rates comparable to WT under both conditions (Figures 6A and 6B). Pollen tube elongation in *slcipk15-1* was similar to WT at 0.1 mM K⁺ but was reduced by approximately 30% at 1 mM K⁺ (Figure 6C). In stark contrast, *slcipk26-1* pollen appeared dehydrated and failed to germinate *in vitro* (Figures 6A and 6B), precluding measurement of tube elongation (Figure 6C).

**Figure 6.**
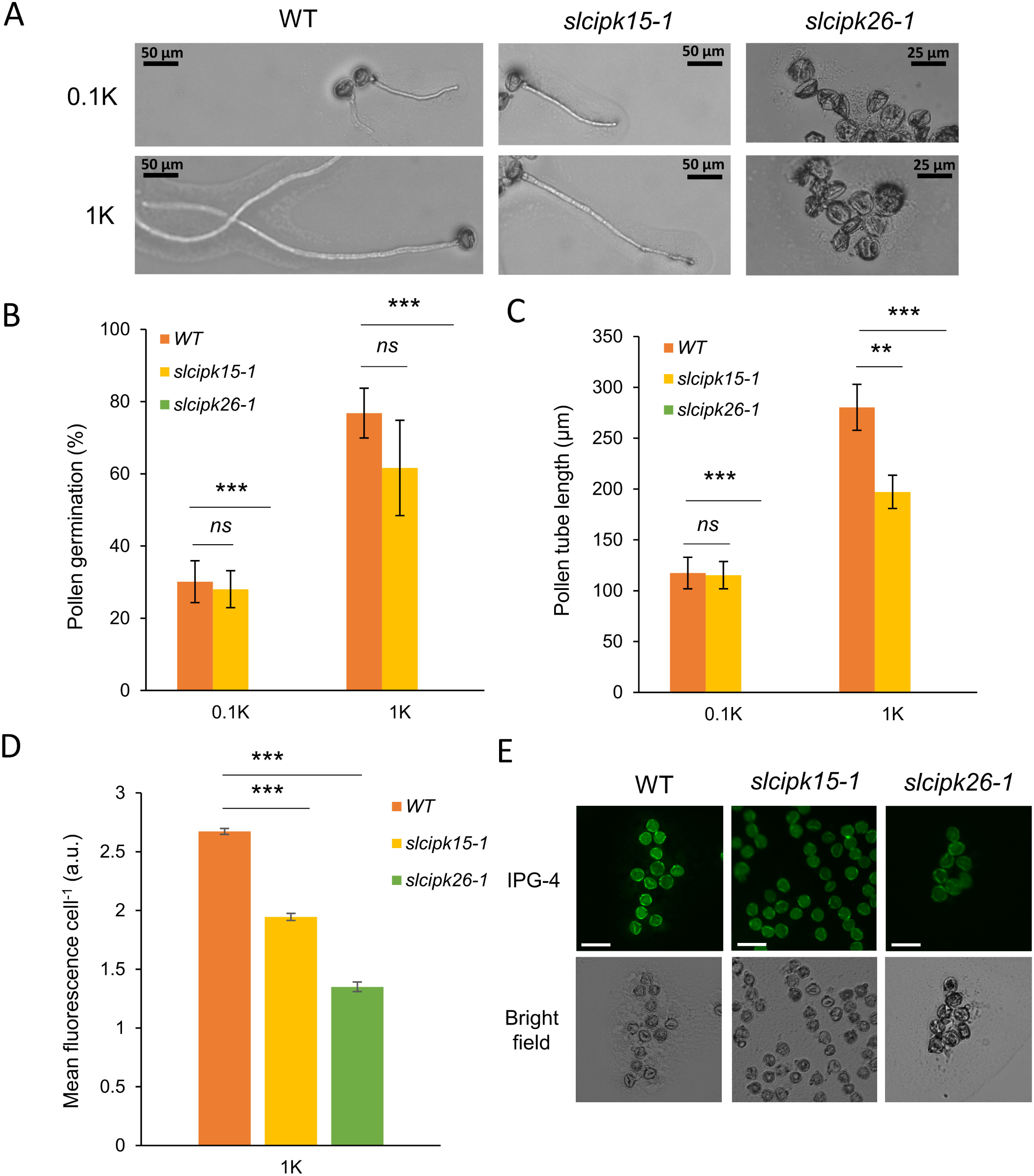
Pollen from *slcipk26* plants showed deficient germination. (a) Representative images of WT, *slcipk15-1*, and *slcipk26-1* pollen incubated in germination media. (b) Germination rate of pollen grains. (c) Pollen tube length. (d) Relative IPG-4 (ION Potassium Green – 4) fluorescence of WT, *slcipk15-1*, *slcipk26-1* pollen grains at 1 mM K^+^ observed under an epifluorescence microscope. (e) Representative images of pollen stained with IPG-4. Scale bar = 50 μm. Data are mean values ± SE (n = 2–7 microscope fields involving 13-78 grains in (b), n = 22-44 tubes in (c), and n = 211-339 grains in (d)) and *, **, and *** indicate p < 0.05, p < 0.01, and p < 0.001, respectively, in Student’s t-test. ns denotes not significant.

Previous studies demonstrated that *slhak5* pollen exhibited reduced K⁺ content, correlating with impaired germination and tube elongation (Nieves-Cordones et al., 2020). To assess K⁺ levels in mutant pollen, the K⁺-sensitive dye IPG-4 was employed. In 1 mM K⁺ medium, *slcipk15-1* and *slcipk26-1* pollen contained 27% and 50% less K⁺ than WT, respectively (Figures 6D and 6E). These K⁺ deficiencies aligned with the observed germination and elongation phenotypes, implicating SlCIPK15 and SlCIPK26 as facilitators of K⁺ uptake in pollen via activation of SlHAK5, with SlCIPK26 playing a more critical role in pollen viability.

### Reduced seed number in *slcipk26* fruits

Given that *slhak5* mutants produce nearly seedless fruits due to reproductive defects (Nieves-Cordones et al., 2020), fruit phenotypes of *slcipk15* and *slcipk26* mutants were characterized. Fruit morphology was similar across genotypes, except *slcipk26-1* fruits contained markedly fewer seeds (Figure 7A). Fruit weight was consistent among genotypes (Figure 7B), while *slcipk26-1* produced slightly fewer fruits per plant compared to WT; *slcipk15-1* did not differ significantly from WT (Figure 7C). WT and *slcipk15-1* fruits yielded 20.5 ± 2.3 and 22.3 ± 1.8 seeds per fruit, respectively (Figure 7D). Despite reduced pollen tube elongation, *slcipk15-1* maintained fertility comparable to WT. Conversely, *slcipk26-1* fruits contained only 2.2 ± 0.9 seeds per fruit, mirroring the near-sterile phenotype of *slhak5* fruits (1.5 ± 1.5 seeds per fruit) (Nieves-Cordones et al., 2020). These results were validated with second mutant alleles (Figure S3D), confirming that *slcipk26* phenocopies *slhak5* fertility defects.

**Figure 7.**
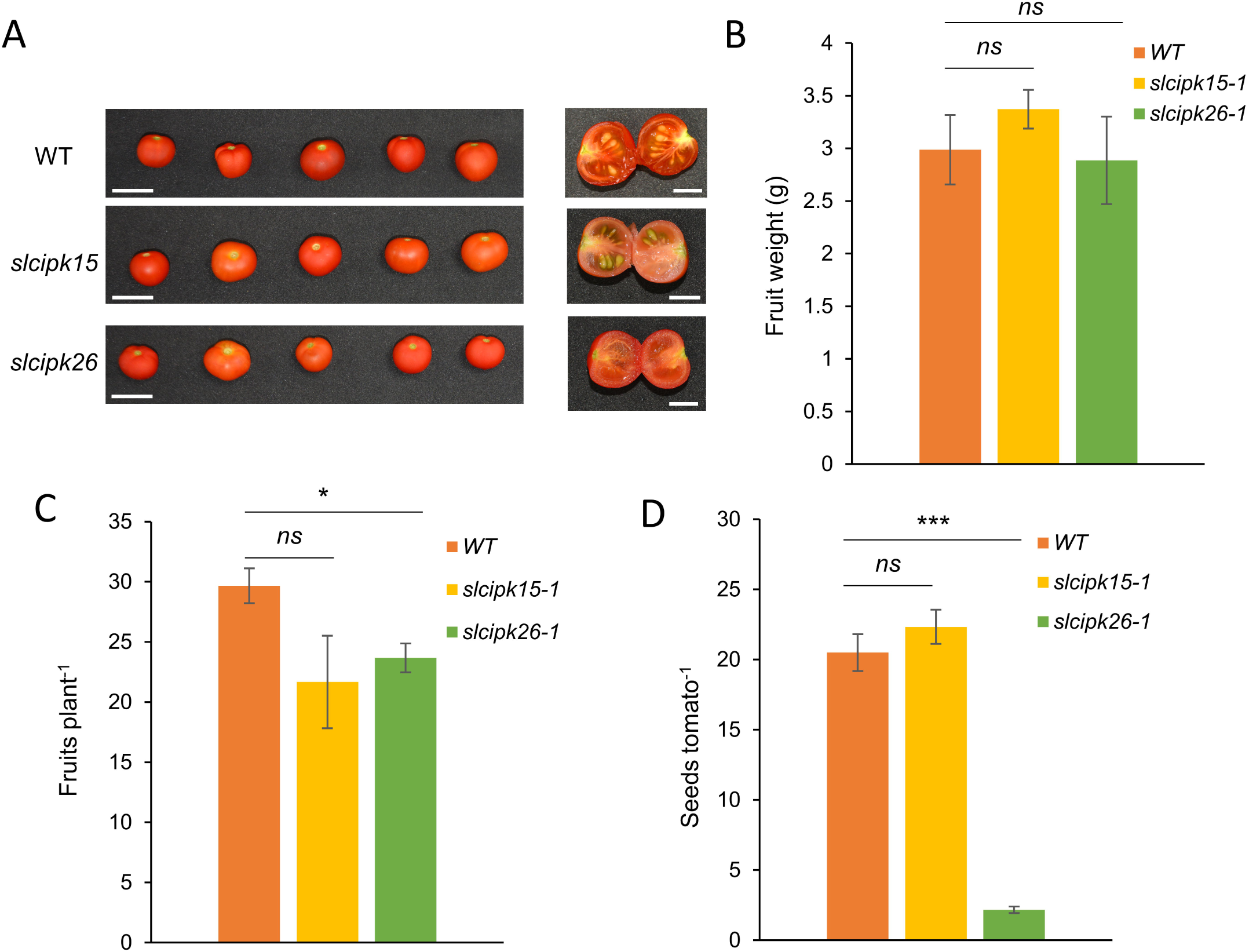
Tomato fruits from *slcipk26* plants had a reduced number of seeds. (a) Photographs of tomato fruits from WT, *slcipk15-1*, and *slcipk26-1* mutant plants. Left panel: Photographs of whole tomatoes. Scale bar: 2 cm. Right panel: Photographs of tomatoes cut into two halves. Scale bar: 1 cm. (b) Fruit weight, (c) fruits per plant, and (d) seeds per plant of WT, *slcipk15-1*, and *slcipk26-1* plants. Data are mean values ± SE (n=6 in (b), n = 3 in (c), and n = 6 in (d)). * and *** indicate p < 0.05 and p < 0.001, respectively, in Student’s t-test. ns denotes not significant.

A recent report noted pistil physiology abnormalities in *slhak5* plants (Jesús Amo et al., 2024). To determine whether *slcipk26* pistils exhibited similar defects, crosses were performed using WT pollen on emasculated *slcipk26-1* plants, with emasculated *slcipk15-1* flowers as controls. Seed counts from 37 *slcipk26-1* crossed flowers yielded 18 fruits with an average of 0.3 ± 0.3 seeds per fruit (Figure 7C). Crosses on *slcipk15-1* pistils produced 21 ± 9 seeds per fruit (n = 3). The *slcipk26* pistil phenotype closely resembled that of *slhak5* pistils crossed with WT pollen, which yielded 0.4 ± 0.1 seeds per fruit (Jesús Amo et al., 2024). These findings implicate SlCIPK26 in pistil physiology, paralleling the role of SlHAK5.

## Discussion

The present study, in conjunction with our previous investigation of AtHAK5 (Lara et al., 2020), provides critical insights into the molecular evolution of the CIPK/CBL/HAK5 signaling network. Notably, only a limited subset of CIPK proteins possess the capacity to activate HAK5 transporters in both tomato and Arabidopsis (Figure 1) (Lara et al., 2020). While certain homologous CIPKs, such as CIPK23, CIPK9 and CIPK26, regulate HAK5 transporters across these species, other HAK5 regulators occupy divergent phylogenetic positions (Figure S2). This disparity underscores that overall protein homology is an unreliable predictor of HAK5 activation, implicating that discrete protein domains may mediate the specific recognition of HAK5 by CIPKs. In Arabidopsis, AtCIPK1, AtCIPK9, and AtCIPK23 activate AtHAK5 both in yeast and *in planta* (Lara et al., 2020), exemplifying the cooperative regulation of root K^+^ uptake by these kinases. Conversely, the tomato SlHAK5 activators identified herein, SlCIPK15 and SlCIPK26, appear uncoupled from root K^+^ uptake (Figures 4C and S3C) and instead associate with reproductive traits (Figures 6 and 7).

The Arabidopsis orthologs of SlCIPK15 and SlCIPK26, namely AtCIPK15 and AtCIPK26, exhibit distinct functional profiles. Unlike SlCIPK15, AtCIPK15 fails to activate AtHAK5 in the presence of AtCBL1 (Lara et al., 2020) and functions as a negative regulator of root NH_4_ ^+^ uptake via phosphorylation of the AMT1;1 transporter (Chen et al., 2020). The AtCIPK26/AtCBL1 complex activates AtHAK5 to a comparable extent as the SlCIPK26/SlCBL9 complex does for SlHAK5 (Figure 1). Beyond its role in AtHAK5 activation, AtCIPK26 modulates reactive oxygen species (ROS) signaling through activation of the NADPH oxidase RBOHF (Drerup et al., 2013) and contributes to vacuolar ionic homeostasis in concert with AtCIPK3/9/23 and AtCBL2/3 (Mogami et al., 2015; Tang et al., 2020b, 2015). The observed overaccumulation of K^+^ in *slcipk26* shoots under +K conditions and in roots under –K conditions (Figures 4B and S3B) may reflect a conserved role for SlCIPK26 in vacuolar K^+^ homeostasis, paralleling the function of its Arabidopsis homolog.

Effective regulation of a target protein by a CIPK/CBL complex necessitates several factors: (1) molecular recognition and functional modulation of the target, (2) spatiotemporal co-expression with the target, (3) ability to outcompete alternative protein regulators for target binding, and (4) capacity to navigate competition among multiple target proteins of the complex. Although *SlCIPK15* and *SlCIPK26* transcripts are detectable in roots (Figure 5), these kinases do not appear to modulate root K^+^ uptake via SlHAK5 (Figures 4C and S3C). By contrast, SlCIPK23 constitutes the principal regulator of SlHAK5 in roots (Amo et al., 2021). The lack of SlHAK5 activation by SlCIPK15 and SlCIPK26 in roots may arise from molecular displacement by SlCIPK23 or preferential affinity of SlCIPK15 and SlCIPK26 for alternative targets. Intriguingly, the *slcipk26* mutant phenocopies the *slhak5* mutant with respect to fertility-related phenotypes, including poor germination, pollen K^+^ accumulation, pistil function, and reduced seed set per fruit (Figures 6, 7, and S3) (Nieves-Cordones et al., 2020). Moreover, *SlCBL9* expression peaks in stamens and is substantial in pistils (Martínez-Martínez et al., 2024), supporting the notion that the SlCIPK26/SlCBL9/SlHAK5 module operates predominantly in reproductive tissues, whereas the SlCIPK23/SlCBL1-9/SlHAK5 complex governs root-specific K^+^ uptake (Amo et al., 2021). Collectively, these findings suggest that distinct CIPK/CBL complexes enable precise tissue-specific regulation of shared target proteins, a phenomenon that, although not widely documented, likely extends across plant species.

The induction of parthenocarpic fruit development holds significant agronomic value for the tomato industry, as such fruits exhibit enhanced qualities for juice extraction and improved organoleptic properties (Varoquaux et al., 2000). Both *slhak5* and *slcipk26* mutants exhibit a drastic reduction in seed number per fruit, averaging approximately one seed per fruit compared to 15–20 in wild-type plants (Nieves-Cordones et al., 2020) (Figures 7D and S3D). This seed paucity stems from compromised fertilization, attributable to defective pollen and pistil functions in both mutants (Figure 6) (Jesus Amo et al., 2024; Nieves-Cordones et al., 2020). Notably, *slcipk26* mutants retain normal K^+^ uptake under low external K^+^ conditions (Figure 4C), a phenotype distinct from *slhak5* mutants, which are impaired in this function (Nieves-Cordones et al., 2020). Furthermore, *slcipk26* plants exhibit largely unaltered growth and development, aside from reduced seed number and a modest decrease in fruit yield relative to wild-type (Figures 7C, 7D, and S3D). These attributes position SlCIPK26 as a promising genome-editing target to produce parthenocarpic tomato with minimal detrimental effects on growth and nutrient uptake.

## Acknowledgements

This work was funded by grant PID2022–137655OB-I00 Ministerio de Ciencia e Innovación, Spain (to FR and MN-C) and Fundación Séneca for grant 22563/PI/24 (to FR). Additional funding was obtained from Consolidation grant CNS2022–135151 (to MN-C), Grant CNS2022–135151 funded by MCIN/AEI/ 10.13039/501100011033 and European Union NextGenerationEU/PRTR (to MN–C). All grants from Spanish ministries were co-financed by the FEDER Fund. AM-M received a FPU predoctoral contract from Ministerio de Educación, Cultura y Deporte, Spain (FPU20/04469). We thank the ionomics and plant transformation services of CEBAS-CSIC for their contribution in this work. The tool Paperpal was used to correct language mistakes and refine text. All suggestions made by this tool were supervised and approved by the authors.

## Supplemental material

**Figure S1.** Production of *slcipk15* and *slcipk26* mutant plants using CRISPR-Cas. (a) *SlCIPK15* locus. (b) *SlCIPK26* locus. Exons are represented by white boxes and introns by black lines. The genomic sequences targeted by the sgRNAs are shown in red. The PAM sequences are underlined. (c) Mutated alleles at sgRNA1 and sgRNA2 target sites in *slcipk15* mutant lines. (d) mutated alleles at sgRNA1 and sgRNA2 target sites in *slcipk26* mutant lines.

**Figure S2.** Phylogenetic tree of tomato and Arabidopsis CIPK proteins. The tree was constructed using the parsimony method in Seaview software. Arabidopsis and tomato proteins are shown in black and red, respectively, in the figure. The number of * denotes the degree of tomato or Arabidopsis HAK5 activation in yeast when co-expressed with the given CIPK together with CBL1 or CBL9 (*** is the maximum degree of HAK5 activation and absence of * is the lowest).

**Figure S3.** Phenotypes of *slcipk15-2* and *slcipk26-2* mutant plants. Tomato plants were grown for 14d in +K solution, then a group of plants was starved of K^+^ for 7 d. On the last day of the experiment, a Rb^+^ uptake assay at 20 µM was performed, and the plants were harvested. (a) Dry weight, (b) K^+^ content, and (c) net Rb^+^ uptake rates of WT, *slcipk15*-2, and *slcipk26*-2 plants grown under +K or –K conditions. (d) Seeds per tomato in WT, *slcipk15*-2, and *slcipk26*-2 plants grown until fructification in pots. Data are shown as boxplots (n = 3-4) in (a), (b), and (c). Data are mean values ± SE in (d) (n=29-39). *, ** and *** indicate p < 0.05, p < 0.01 and p < 0.001 in Student’s t test, respectively.

**Figure S4.** Expression in different plant tissues of *SlCIPK15* and *SlCIPK26* transcripts according to the Solgenomics database (https://solgenomics.net/).

**Table S1.** Coding sequences of SlCIPK proteins used for expression in yeast.

**Table S2.** Primers used in this study.

