## Supplemental files for "SlCIPK26 enhances tomato fertility by activating the K^+^ transporter SlHAK5 in reproductive tissues"

A

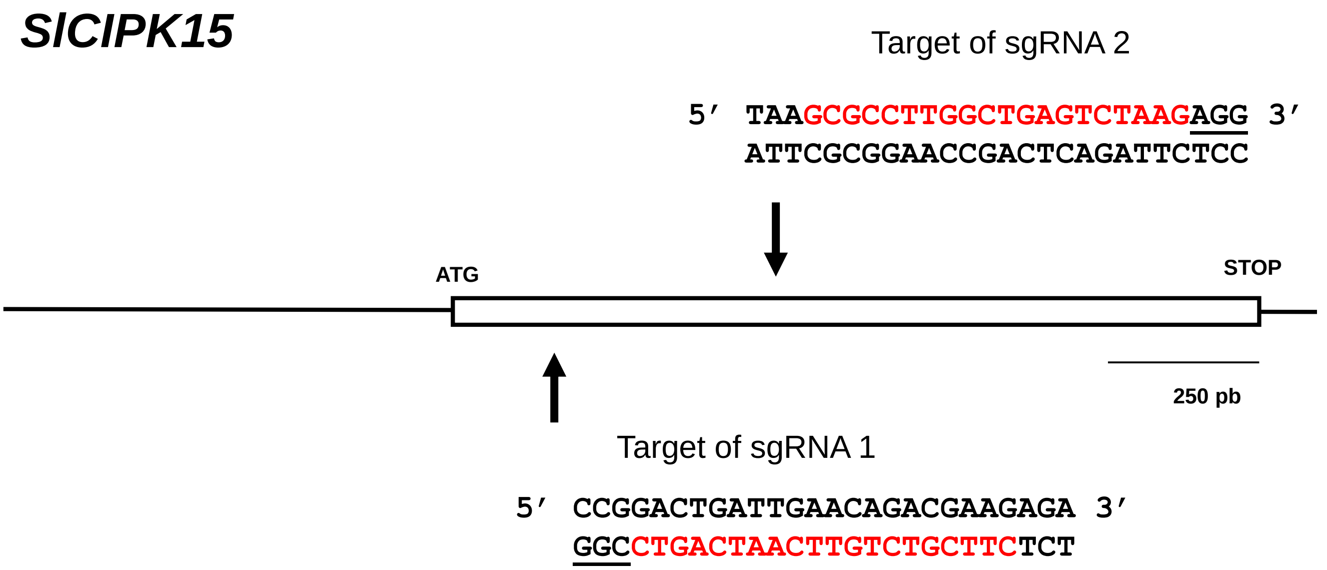


B

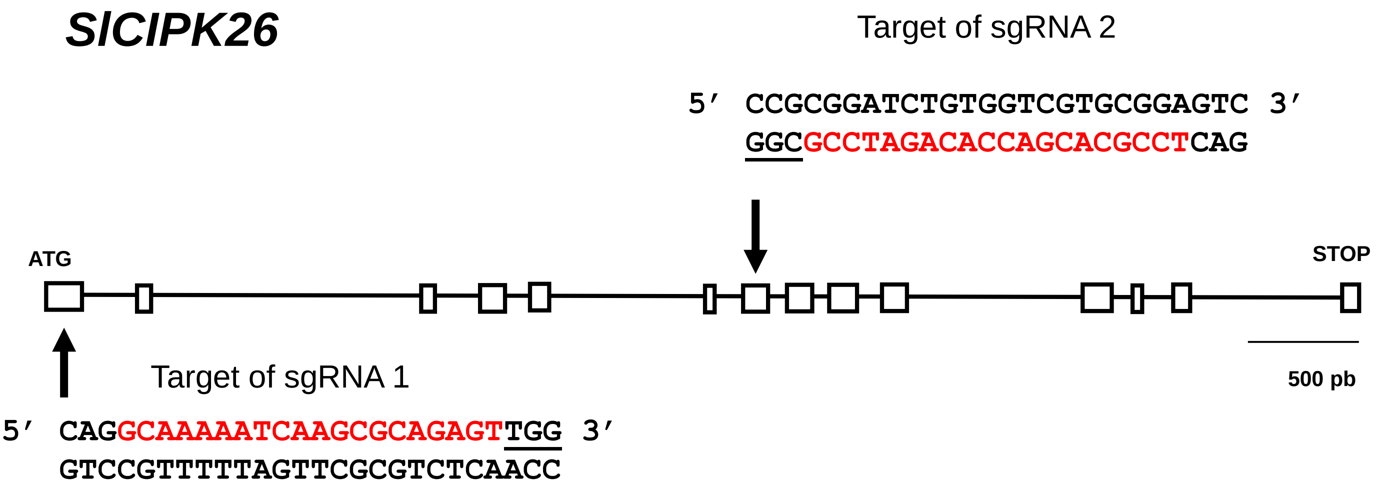


C

| Name | Strand | Target of sgRNA1 | Genotype |
|---|---|---|---|
| SlCIPK15 (WT) | - | 5´ TCTCTTCGTCTGTTCAATCAGTCCGG 3′ | |
| *slcipk15-1* *slcipk15-2* | - | 5´ TCTCTTCGTCTGTTCAATCAGTCCGG 3′ | Non-mutated |

| Name | Strand | Target of sgRNA2 | Genotype |
|---|---|---|---|
| SlCIPK15 (WT) | + | 5′ TAAGCGCCTTGGCTGAGTCTA--AGAGG 3′ | |
| *slcipk15-1* | + | 5′ TAAGCGCCTTGGCTGAGTCTA----AGG 3′ -2 bp | Homozygotic |
| *slcipk15-2* | + | 5′ TAAGCGCCTTGGCTGAGTCT---AGAGG 3′ -1 bp<br>5′ TAAGCGCCTTGGCTGAGTCTAagAGAGG 3′ +2 bp | Bi-allelic |

D

| Name | Strand | Target of sgRNA1 | Genotype |
|---|---|---|---|
| SlCIPK26 (WT) | + | 5´ CAGGCAAAAATCAAGCGCAGAGTTGG 3′ | |
| *slcipk26-1* *slcipk26-2* | + | 5´ CAGGCAAAAATCAAGCGCAGAGTTGG 3′ | Non-mutated |

| Name | Strand | Sitio diana sgRNA2 | Genotype |
|---|---|---|---|
| SlCIPK26 (WT) | - | 5´ GACTCCGCACGACCACAGAT-CCGCGG 3′ | |
| *slcipk26-1* | - | 5′ GACTCCGCACGACCACAGATtCCGCGG 3′ +1 bp<br>5′ GACTCCGCACGACCACAGA---CGCGG 3′ -2 bp | Bi-allelic |
| *slcipk26-2* | - | 5′ GACTCCGCACGACCAC-----CCGCGG 3′ -4 bp<br>5´ GACTCCGCACGACCACAGAT--CGCGG 3′ -1 bp | Bi-allelic |

Figure S1

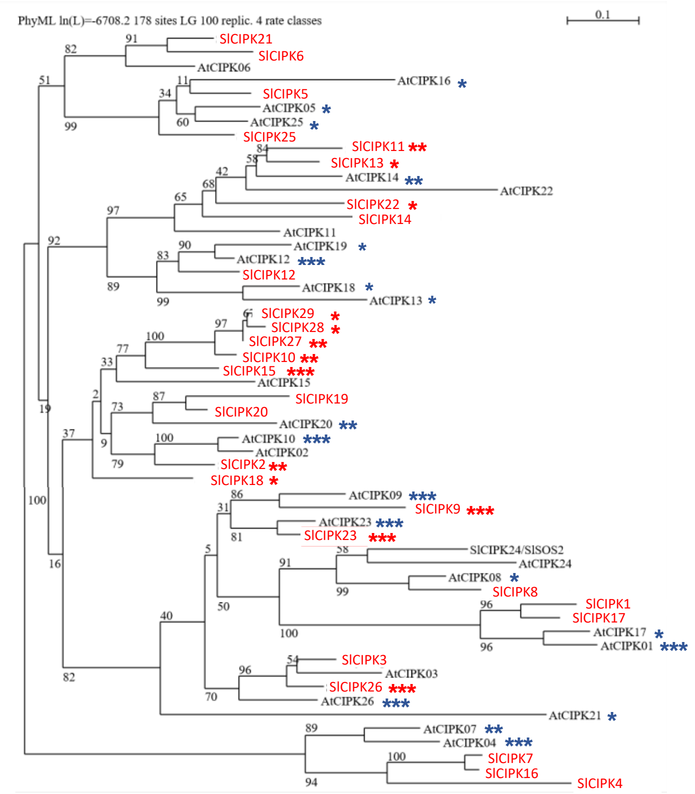
PhyML ln(L)=-6708.2 178 sites LG 100 replic. 4 rate classes
0.1
SlCIPK21
SlCIPK6
AtCIPK06
AtCIPK16 *
SlCIPK5
AtCIPK05 *
AtCIPK25 *
SlCIPK25
SlCIPK11 **
SlCIPK13 *
AtCIPK14 **
AtCIPK22
SlCIPK22 *
SlCIPK14
AtCIPK11
AtCIPK19 *
AtCIPK12 ***
SlCIPK12
AtCIPK18 *
AtCIPK13 *
SlCIPK29 *
SlCIPK28 *
SlCIPK27 **
SlCIPK10 **
SlCIPK15 ***
AtCIPK15
SlCIPK19
SlCIPK20
AtCIPK20 **
AtCIPK10 ***
AtCIPK02
SlCIPK2 **
SlCIPK18 *
AtCIPK09 ***
SlCIPK9 ***
AtCIPK23 ***
SlCIPK23 ***
SlCIPK24/SlSOS2
AtCIPK24
AtCIPK08 *
SlCIPK8
SlCIPK1
SlCIPK17
AtCIPK17 *
AtCIPK01 ***
SlCIPK3
AtCIPK03
SlCIPK26 ***
AtCIPK26 ***
AtCIPK21 *
AtCIPK07 **
AtCIPK04 ***
SlCIPK7
SlCIPK16
SlCIPK4

**Figure S2**

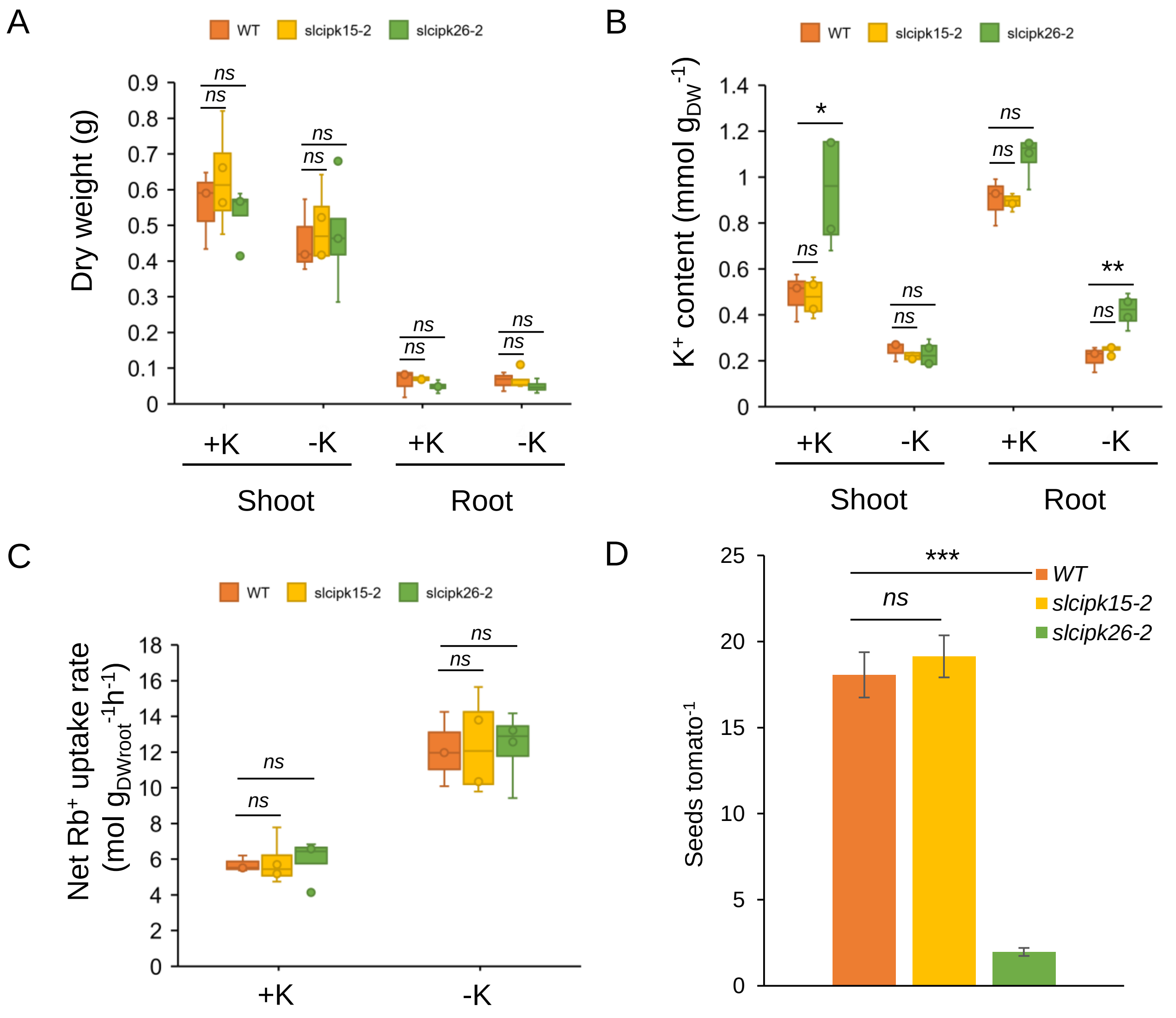
A
WT
slcipk15-2
slcipk26-2
Dry weight (g)
0.9
0.8
0.7
0.6
0.5
0.4
0.3
0.2
0.1
0
ns
ns
+K
-K
+K
-K
Shoot
Root
B
K+ content (mmol gDW-1)
1.4
1.2
1
0.8
0.6
0.4
0.2
0
*
**
ns
C
Net Rb+ uptake rate
(mol gDWroot-1h-1)
18
16
14
12
10
8
6
4
2
0
D
***
WT
slcipk15-2
slcipk26-2
Seeds tomato-1
25
20
15
10
5
0

Figure S3

A

*SlCIPK15*

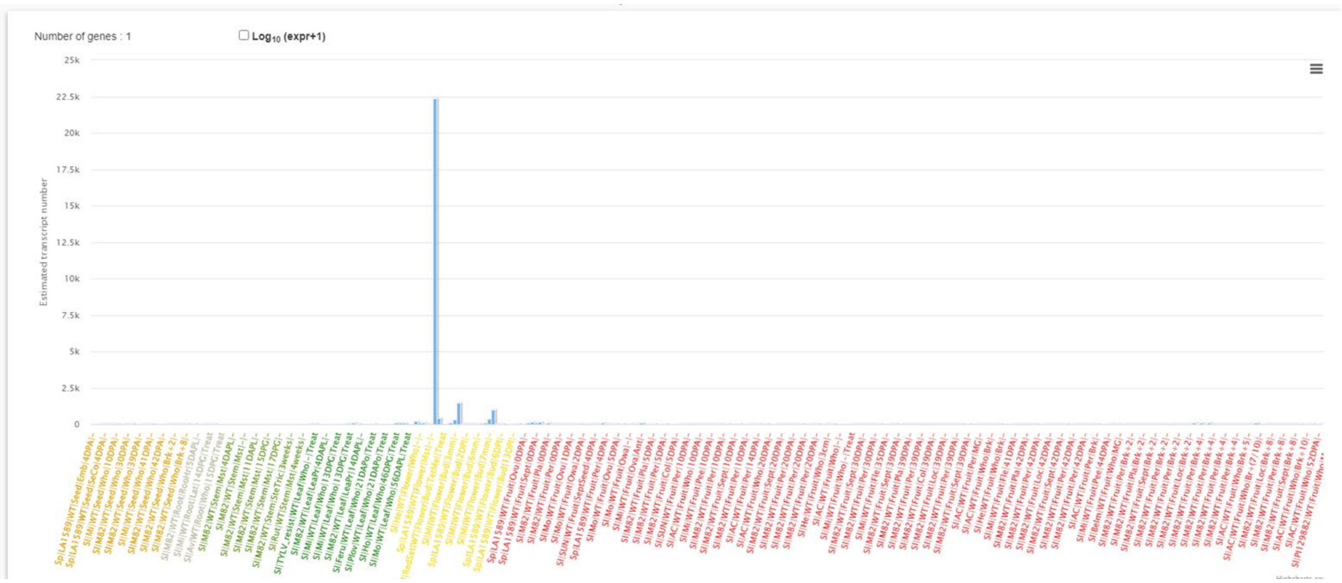


B

*SlCIPK26*

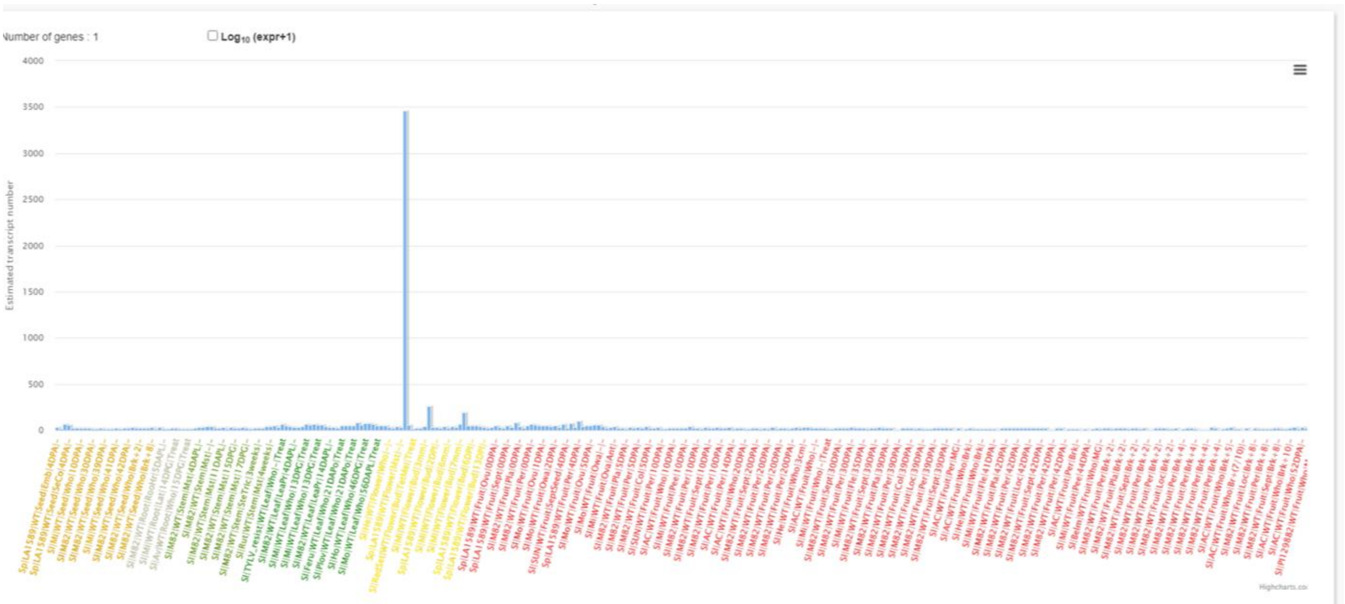


Figure S4

**Table S1.** Coding sequences of SlCIPK proteins used for expression in yeast.

| SlCIPK number | Locus | ORF sequence |
|---|---|---|
| **SlCIPK1** | Solyc05g053210 | atggtgttggtacaacaggaagacgaaataagggtgggtgcagggaagaagggaatgaggctagggaaatatgaagtggggaagacacttggtgaaggaaattttggtaaagtgaagtatgctagacatgtagagactggtcaatctttcgccattaaaattttggagaagagtcggatccttgacctcaagtctactgatcagatcaagagggagattggaactttaaaactcctcaaacatcctaacgttgtccgattatacgaggtcttagcgagcaaaagcaagattttcatggtgctagaatacgtgaatggggtgaattattcgacagaatcgtttccaaaggaaaactctcagaggcccaaggtaggaagctcttccaacaattggttgatggcgtaagttactgtcacgacaaaggtgtcttccacagagatctcaagctagaaaacgtcctcattgatttgaggggaaacataaaaataacggactttgggctcagtgcgttaccccaacactttagggatgacggattgttgcacaccacttgcggtagtccaaactatgtcgctccagaaattctttctaacagaggatatgatggcgcggcatctgatacctggtcatgtggtgttatcttatatgttattctcactggttatttaccctttgatgatagaaatcttgccgtactttatcaaaagatacttaagggggaagttcatataccaaaatggctatctgcaggagcaaagaacctcataaagaggattcttgatcccaacccgcatactcgtataacaatggcacaaatcaaagaagatgcatggtttaaacaagactatacacctgtaaacactgatgatgaagacttggaaagtgatgatcacgtctgcaccgtgcatgaaacggatgatcacgtctgtaccgtgcatgaattgccacttgatgctcaaagagatcccgaatcgccttgccttatcaatgcctttgaacttataggaatgtcatcatgtctggatctttctggatttttgagaaagaggatgtttccgagaggaagatcagatttacatccagtctctctccaaaacagttgttggagaggattgagaatatggtgacacagatgggatttcatgtccagaaaagacacggaaggttgaaagtgatgcaagagcagaaaggccacaaaaacccggctagtctattggtagttgctgaggtcttcgagattagcccatccttgtatgttgtagagttacaaaagtcttcaggggattcaacagtatatagacagatgtgtaataggttatcgaatgatttgggagtccaccgtaatgaggagctcctacctactgtttgttgtgatagctag |
| **SlCIPK2** | Solyc06g007430 | atggccaataaaggaagcatactgatggagcggtatgaagtgggcagattattaggtcaaggtacatttgctaaggtttactatgcgaggaatatcaaaaccggacagagtgttgccatcaaagtcatagacaaggaaaaagttgtcagggtcgggctcatgaatcagatcaaacgggagatatctgttatgaaactagtcagacatccaaatattgtgcatctttacgaggtcatggcgacaaaaaccaagatatactttatcatggagtattgtaaaggaggtgagctctttaacaaggtagctaagggaaggctaaaagaggacgcagcacggaaatattttcagcagttgataaatgctgtagatttctgccatagtagggggtgtctatcaccgggatttgaaacctgaaaacttgctgttagatgacgatgaaaacttaaaaatttcagattttggtttaagtgctctcgttgagtcaaagcaccaagacggactcctccacacgacgtgtgggactccagcttatgttgctccagaggtgattaacagaagaggctatgatgggactaaggctgatatctggtcgtgtggggttgtcctatacgttctgttggctggttatcttccatttcaagactcaaatttgatggaaatgtataggaagattgggaaagctgattacaaatgtccgagttggtttccaccagaagcccgacgtctactttcaagaatgttggatcctaatccaagttcaagaatttcccttgcaaaaattagagcaagttcgtggttccggaggggaatttcaacttcctctaaatctacagtagtagacgacgtaagcacggatttagcttcagcaaataaagaggacaagcaagaagtggcttcgattccaaagttgaatgcttttgatatcatttccctttatgctaagtttgatttatccagattgttcgaggaaccttgtttaaagaaagaaactaaattcacatcccggaaaactgcatcagtgatcatatccaagctcgaaaatattgctaaacatctgaagctgaaagtttgcaaaagggatgcaggattactgaagttcgaaggaacaaaagaaggaaaaaagggaactttgagcattgacgtggagatctttgaggttgttgaagcttttcatttggtggaagttaaaaaatcaaacggggatacattggaatatcaacagatattgaacgaaggcctaagaccgggtcttcaagatatcgtttggacttggcaagaagatcaacagctgcctcagcagtcggaagaccagcttcacgagcagacaccaaataatcaacttcagcaggaacaacatttagataatcagcaacaacctccggagcagcagcaactgctactccaaaaccatctgatacagcaagagcagttaccttga |
| **SlCIPK3** | Solyc01g008850 | atgaatcggaccaaaatcaagcgtagagttggtaaatatgaagttggaaggacaattggtgagggaacgtttgcgaaagtcaagtttgcaaagaattcagagacgggagaaagtgtggcgatcaagatcctcgataaagataaggtccttaagcacaaaatggctgaacagataaagcgggaaatagctacaatgaagttgatcagacatcctcatgttgtacagttatacgaggttatgggaagcaagacgaagatatttattgttttggagttcgttacaggtggagagttattcgataaaattgtaaatcatggacgaatgcgtgaagaagaggcaaggaagtattttcagcaacttattcatacggttgattattgccatagtagaggagtctatcaccgagatctaaagcctgaaaatttactgttggattcttcgggtgaccttaaggtatctgattttggattgagtgctctatcccagcaagtcagggctgatggcttactgcataccacctgcggaactcctaactatgttgcacctgaggtactgaatgatcgaggatacgatggggcaacagcagacctctggtcgtgcggagtcatactctttgtactgcttgcaggttacttgccttttgatgactccaatcttatgaacctctataacaaaatctctgctgctgaatttacgtgcccaccttggatctcttttggtgctatcaagttgattactcgcatcttggatccgaatcctacgacacgtattaccgtgcctgaaattttggaggatgaatggtttaagaaagattatagaccacctgttttcgatgaaatagaagatgcaaacctgga |

| | | |
|---|---|---|
| | | tgacgttgaagcagtcttcagggactctgaagaatatcacgtaacggaaagaagagaagaga agccaacttccatgaatgctttcgagttgatttctatgtcacaaggtcttaaccttggcaatctcttcg acgaacagggattcaagagagagacaaggttcacgtctaaatgtccggctaatgagataatca gtaagattgaagaagctgctaagcctctcggttttgacgttcacaaaaagaactacaagatgag gcttcaaaatctgaaagccggtagaaaagggaaccttaacgtttccactgaggtgtttcaagttg ctccttctcttcatatggtcgaggtgcggaagtcaaaaggagacactttggaattccacaagtttta caagaatctttcaacctgcctagataatgttgtatggaaaactgaacaagacatggaagaaaa aaaatga |
| **SlCIPK4** | Solyc03g005330 | atggaggcgaaatgcactcaaccaccctcctccgccacgataatcagacgaaccaccagca ccaacagaagtggttccggaaacggaaccatcattctcggcaagtaccaacttggccgtctatt aggccgtggtagctttgctaaagtctaccacggcctttgtttagacgataacacaaatatcgctatc aaggttatcgataaaaaatcgatcattggcacgaatgtctcaatggagccccgaatcctacgtg aaatctccatcatgtcctgtcttaatcatcccaacataataaaactccatgaagtaatggctacaa aatccaaaatctatcttgtcatggagatcgcctgcagaggcgatcttcatgccaagctaatccgtc atggacggttttctgagtccaccgctcgtttctacttccaccaattagtctccgctctccattactgcc atcaaaatggcgttactcaccgcgacatcaaacctcaaaacattcttcttgatcaaaataacaac atcaaaatctccgatttcgggctctccgccttgcctgaacagctgaaaaacagtctccttcataca gcttgcggtacaccaggttatacagctccagaggtagcttacggaaagggatacaatggcgaa aaggcggattcatggtcttgtggggtaattctatttgcatttctatcgggatatattccatttgattctgg caatttatcaaacatgtaccgtgcaatacaccgtcgcgattttcgattccctaattgggtttcgaaat cagctcggagtgtgatcaacaagttactcgatccaaatccgagtactagattaagtattgagcaa ctaatgaacctttcctggtttaagaaatcgaatcaacaacaaagatcaagcaattgtgtgttcgag aagaatcgtacaaatttgggcgggataaatgcatttgacatattatcaatgtcatcagggttgaatt tatcgaggttattcgagagcgatttgatcaatagagagatgaagttcacgacgaatgctcgaatc gaggaggttgaagagaaggtggtgaaaattggagaaggaggagggtatagagttgaaaga gggaaaggaaggggaattgagttggttaaagggagagttgttttaatggtggaaattttggaggt ggcaatggagttattgttggttgaagttaaggttgttaataatggaggattggaatttgatgattctca atgggaggatttgaaatttgggatgaaggatatttgttag |
| **SlCIPK5** | Solyc08g067310 | atggaggagagacaattgttgtttgacaagtatgaaatcggaaagttattaggtaaagggactttt gcgaaagtgtactatggtaaagaagtggaaacaggggaaagtgtagccattaaagtgataaa aaaagatcaagttcaaaaagaaggaatgatggagcagatcacaagagaaatctcagtgatg gggctggttcgacatccgaatatagttgaactcaaagaagtcatggcgacaaagtctaaaatctt cttcataatggagtatgttaaagggggtgagcttttcgcaaaattagtatccaaaggcaaatttaa agaagatacagcaagaaagtactttcaacagttgataagtgctgttgatttctgtcatagtcgagg tgtttatcatcgcgatttgaagcctgagaacttactccttgatgagaatgaggacttgaagatatctg atttcgggctatcggctttgcctgatgagcagttgagtagaaatgatggtttgcttcatacacagtgt gggacaccagcatatgttgctcctgaggttttaaggagaaaaggttatgatggtgcaaaggctg atatctggtcatgtggggttatcttgtatgtgcttctagctggttacttgccatttcaagatgagaatgt gatgaacatgtacaagaagatcttcaaggcgtatttcgagttccctccttggttttcaatggattcga ggcgtttgatatctaaacttttaatggctgatccagacaggaggatcactattcaaggtatcatgag ggttccatggtttaggaaggatttcgccatgccacgggctttttcaatccaagattttcaaaagttag aaaatgatcatgttgaagaaggaatgagcagcaaacaaggaggaggggtcctaaggaaatc cccttcatcccctgcatttttcaatgcattcgagttgatctcatccatgtcttctggtttcgacttgtctag cttgtttgagatcaaagggaaggcatcatccatgttcacttcaaggagcactgcccgggatgttat ccgtaaaatggagaaaatggctaaagttatgaggtataaggtttatagggtaaaaccattcaagt taaagatgcaatgccccgaggaaggacgaaaagggcggttgttggtgaccgctgaggtgttca aggtggccccggaagtcaccgtggtcgagctctctaagtcctccggcgacaccttggagtataa caagttttgtgaagaggaagttaggcctgcgttgaaggacattgtttggacatggcaagggaca ggtactggaaatgctgatattgataacaaacaggacttggagcaataa |
| **SlCIPK6** | Solyc07g005440 | atggcccctgaagagaaatgtggagctttgaacggaaaatatgaactcggtcgacttttaggcc atggaacattcgcgaaagtttatcacgctcgaaacgtgaagaatggtaaaaatgttgccatgaa agtagtggggaaagaaaaagttattaaagtaggtatgatggatcagatcaagcgagaaatctc ggttatgaaaatggtgaaaaacccaaacatagtcgagctccacgaagtcatggcgagtaaaa caaagatttacttcgccatggagttcgttaaaggaggtgagcttttcgcgaagatcgcgaaaggc aaggtgagagaggatgttgctagaggttacttccagcaactgatctcagctatcgacttctgtcat agcagaggagtatttcatcgcgatttgaaacctgaaaacttattgttggatgaagaaggaaatct caaaattactgattttgggctaagtgcatttacagaacatcaaagacaagacggattgcttcatac aacttgcggaactccggcttatgttgcccctgaaattattggtaaaaaaggctacgatggttcaaa agcagacatatggtcatgtggagtgattctctatgttttactagctggtttttaccatttcaagacgaa aacatcatggcgatgtataagaagatttaccggggtgatttcaaatgtccaccttggttttcatctga agctaggaggttgatcacgaagatgttggatccgaatcctaattcaagaatcactacttcaaaga tcatggattcgtcatggttcaaaaaatcgatcccaaagacgttaaggaacaaagatgaggaag |

| | | |
|---|---|---|
| | | aatttgcatttgcatttgcatcagataagtctagtaaacaagttgagacaatgaacgcgtttcatatc<br>atttctttatctgaagggtttgatttgtcaccattgttcgaagagaataaaaggaatgagaaagaac<br>agatgagattcgcgactacaatgtcagcaagtagtgtgatatctaagcttgaggaggtagcaaa<br>gacaacgaatttcatcgtaaaaaagagtgattcctgtgtaaaactacaaggacaagtggtggg<br>gagaaaagggaaattaggaattgctgctgatatatttgctgtgacaaattcatttctagttgtggaa<br>gtgaataaagcaagtggtgatacattggagtataatcaattctgcagcaaagagctaagacca<br>gcacttaaggatattgtttggacatcagcaacatag |
| **SlCIPK7** | Solyc09g083090 | atggatgtaaaacaaccccaaccacctccatttaccgtcaaaatcagaagaaccaccagttcc<br>gatggtagtggatctggatctatcattcttgggaaataccaattggttcgtctgttgggtcgaggtag<br>ctttgctaaagtgtaccttggccgttgtttagacgataacactgaagttgctgttaaagttatagata<br>aatccagtactgccattgatgcttctatggagccacggattatccgtgaagtctccgccatgcgcc<br>gccttaatcaccacccgaatatcctcgaactttttgaagttatggcgacgaaaactaagatctattt<br>cgttatggaactagctcacggcggtgagcttttcacaaagcttaatcgccgtggccggttttctgaa<br>tccaccgccagattttatttccatcatcttgtctctgctttacatttctgccaccaaaacggcgttgctc<br>atcgtgatatcaagccacaaaatctactcctagacaaagagggtcatctcaaaatctccgatttc<br>ggactctccgccttgccggagcagttacagaacggtctcctttatactgcgtgtggtacgccggc<br>gtatactgctccggaggtagtttacagaagagggtacgatggtgctaaggcggatgcttggtcat<br>gtggggttattctctttgtgttcctcgccggaagcttaccgttcgatgatagcaatttgcctaacatgg<br>ttaaagctatacaccggcgtgaatataagtttcccgattgggtttcaaagtcagctcagaggataa<br>ttaaccggctgcttgatcctaatcctgaaacgagatacggaattgtagagctcatgaacactccat<br>ggtttaagaaatcatcgtcgatgaaaccagagcaaagcacgaagcgatttggtgaggggatttt<br>ggagaaggaaagcaaacaaatggagagtataaatgcatttgatttaattttaatgtgttcagggtt<br>ggatttatcgtcgatatttgaagaagaattgaacaagaaggagatgagatttacgacaaatgta<br>gaagttaaggtgatagaagagaaggtgatgaatgttggtataaatgcaggatacagagtagag<br>aaaagaaagaatggtggaattgggttggtgaaagggagaagtgttttgttagttgaaatcttgga<br>gttggcaaaagagttgttgttggtggagttcaaagttgttaatggaggatcagaattcgaggatcg<br>tcaatgggaagaattgaaagctggattgaaagaagtagttgtttcatggtag |
| **SlCIPK8** | Solyc04g076810 | ATGGTGGTAAGGAAAGTTGGTAAGTATGAAGTTGGAAGGACAAT<br>TGGAGAAGGAACATTTGCTAAGGTTAAATTTGCTCAGAATACTGA<br>GACAGGTGAAAGTGTCGCCATGAAAGTCCTCGATCGAAGCACTA<br>TCATCAAGCACAAGATGGTTGACCAGATAAAGCAGGAGATATCC<br>ATAATGAAGCTTGTTAGACATCCATATGTAGTTCGATTACATGAG<br>GTTATAGCAACTCGCACGAAGATCTATATTATCTTGGAATTTATCA<br>CAGGCGGGGAACTTTTTGATAAGATAGTCCACCATGGACGATTA<br>AGTGAGGCCGAGTCTCGAAGATACTTTCAACAATTGATTGATGGA<br>GTTGATTATTGTCACATCAAGGGAGTTTATCACAGAGACCTAAAG<br>CCTGAAAATCTTCTGCTAGATTCCCAAGCAAATCTGAAAATATCA<br>GATTTTGGACTTAGTGCATCACCTGGCGAAGGAGTCAACATTCTT<br>AAGACTACATGTGGAACTCCCAACTATGTTGCACCAGAGGTTCTT<br>AGTCACAAAGGTTATGATGGTGCTGTGGCTGATATCTGGTCCTG<br>TGGTGTCATCCTTTATGTTCTGATGGCAGGTTATCTCCCTTTTGA<br>TGAGGTTGATCTCACTACACTGTACGCAAAGATTGACAAAGCAGA<br>TTTTTCCTGCCCATCTTGGTTTCCTGTTGGAGCAAAATCTCTGAT<br>ACATCGAATTTTAGACCCAAATCCTCAAACTCGTATTCGGATTGA<br>AGAGATCCGTAATGATGAGTGGTTTAAAAAAAATTATGATCCTGT<br>CAAAGTCATGGAGTATGAAGATGTCAATTTAGATGATATTAATGC<br>AGCTTTTGATGATACTGAGGAGGAAGCATCCAACGAGCAATGTG<br>ACAATGCGGATGCTGGGCCTCTGGCTTTAAATGCCTTTGACCTA<br>ATTATTCTCTCTCAAGGATTGAACTTATCCATATTGTTTGACCGTG<br>GGCAGGACTCAATGAAGCATCATCAAACACGCTTCCTAACACAG<br>AAACCAGCAAAAGTTGTTTTATCAAGTATGGAAGTTGTGGCCCAG<br>TCCATGGGTTTCAAGACCCATATCCGCAATTTTAAGATGAGGGTA<br>GAAGGTCTCTCCACAAACAAGACTTCACATTTCTCTGTAATACTG<br>GAGGTTTTCGAAGTTGCTCCTACATTTTTCATGGTAGACGTTCAG<br>AAAGCAGCCGGTGATGCTAGTGAATTCCTCAAGTTTTACAAGAAC<br>TTTTGTGGCAATCTTGAGGACATTATCTGGAGGCCACCGGATGA<br>ATCATGCAAATCAAAAGTTACAAAAGCAAGGAGTAGAAAGAGATG<br>A |
| **SlCIPK10** | Solyc06g050300 | ATGGTGAAAAATGGAAATGTAGTGATGCAAAAATATGAATTGGGG<br>AGATTATTAGGTCAAGGCAACTTTGGTAAGGTTTATTATGGAAGG<br>GATCTGGAAAGCGGACAAACTGTAGCCATTAAAGTAATTGATAAA<br>GAGAAGGTTCAGAAAGCTGAATTGACTGAACAGACGAAACGAGA |

| | | |
|---|---|---|
| | | GATATCCGTTATGGCAATGGTCAAACATCGACATGTTGTGCAGCT<br>ATACGAGGTCATGGCAACCAAGAGTAAGATTTACTTTGTGATCGA<br>ACATGCCAAAGGTGGCGAGCTTTTCAACAAACTGACAAAGGGGA<br>GGCTCACAGAAGATGTTGCTAGAAAGTTGTTTCAGCAACTGATCA<br>ATGCAGTTGAATTTTGCCACAGCCGAGGTGTTTATCACCGTGATC<br>TCAAACCGGAAAATCTCCTACTGGATGAGAATGGAAACCTAAAG<br>GTCTCAGACTTTGGTTTGAGTGCATTAGCAGAGTCTAAGCGACAA<br>GATGGGTTACTCCACACAACCTGTGGTACACCAGCATATGTTGC<br>TCCCGAGGTGATTGGTAGAAAAGGATATGAGGGTGCTAAAGCTG<br>ACATCTGGTCTTGTGGGGTGATTTTATTTGTCTTGTTGGCTGGTT<br>ATCTTCCATTCTATGACTTAAATCTTATGAATCTGTATAGGAAGAT<br>ATGCAGGGCAGAGTACAAATGCCCTAATTGGTTCCCTCTAGAAG<br>TGCGTAAACTTCTCTCTAGGATCTTCGACCCAAACCCTCGTAAAA<br>GGATTTCAATCGCCAAAATTAAGGAAAGCTCCTGGTTTAAGAAAG<br>GATTGGAATCTAGACATGTGGGAACTAAACAAGTAGTGAACCAAA<br>ATGTTATTGCAGATGGTAATGCTGTTTCCAGTTCAAATTCGGAGA<br>ATAGTACCTCTTCCTCCGACACCAAATTAGAGTTGATAAAACCTG<br>CAATCTTTAGTGCATTTAATATCCTCTGCCAGTTTAATTTGTCTGG<br>TTTATTCATAAACAATGATCAAAAGGAGGAGCTGCTTTTTACATCA<br>GTGGAACCTGTCCCAGTCATCATATCCAAACTTGTGGAAGTTGG<br>CAGGAGTCTGAACCTTGAAGTAAAGAAGAAAGAAGGTGGATTTC<br>TTATATTAGAGGGATTAAATGAGAGCAGATATGAGACCCTGTGCA<br>TTGGCGTGCAAATCTTTGAAATTTCCGTATCCAATTACTTCATTGA<br>ACTGAGTAGGTCAAGTGGTGATGTGATTGATTACCAAAACATGTT<br>GACGCAAACTATCAGACCAGCTCTTGAGGAAATTGTTCTGGCTT<br>GGCTAGGAGTGCAATCTTATCAATAG |
| **SlCIPK11** | Solyc10g085450 | atgccagagatcttacgtgaggagagcagttcatcttcatcaaccaccgccggcaccatcgccg<br>gcgccggcaccggagatttcaagcttaacatatttgggaaatatgaggtcgggaagcttctgggt<br>tgtggtgcttttgcaaaggtttatcatgctagggatgtgaggacgagacagagcgtggcgattaa<br>gacggtaagcaagcagaagatactgaaaggtggtttcacggagcatgttaagagggagatttc<br>tatcatgcgcaggttgcgtcatcctcatatcgtgcgccttcatgaggtgctcgcgacgaagacga<br>aggtctattacgttatggaattcgctaaaggtggagagctgtttacgaaagttagcaaagggaga<br>ttcagcgaggatctcagccggagatattttcagcagttgatctccgccgttgattattgtcattctcgt<br>ggtgtgtatcatcgagatctgaaactggagaatctgttgctcgatgagaattgggatctcaaggtc<br>accgattttgggctcagtgcagtgacggatcagatccgacccgatggattcctccatacgttatgc<br>ggcactccggcatatgtagctcctgaaatcctcgagaagaaaggatacaacggagctaaggt<br>ggatatatggtcatgcggcatcatcctttttgtgctcaacgccggttatttaccatttaccgatgcaaa<br>tctaatggcgatgtatcggaaaatctacaaaggcgaattccgttgccctaaatggacctcacacg<br>gattgaaactgttgcttactcgactcctcgacacgaatcccgacactcgaatctccattgaacaa<br>atcagaaacgatccgtggtttcggaaaggttacaaagaggtgaaatcgcatttcggtgatgaatt<br>tgagttgaaaagtagctccgatttgaatcgcgacggcaagtttctaaatgcatttcatataatctcgt<br>tatcctcaggtgttaatctgtctggattgttgaaatctctaggagggaaagagacagtggatgttga<br>aagatttgtttcagcatcgtcagcggagagaataacacagaggattgaagagattgcaaaggc<br>tgaagggatgagagtcaccggtaaaaatggcgccggtgtaagagttgaggggcaaaatggg<br>aaatttgttttagcgacagagattaaccggttaacggagaaagtggtaatagtagaagttaaacg<br>aaaggagataggagctggagctgatggagaaatatggaaactcaaattcaagccgcagttaa<br>gtgaattctttttgtga |
| **SlCIPK12** | Solyc02g072530 | atggccggcatcgctaccgccggcgacaccagcaccagtaccagcagcaacctagcgccg<br>acgaggagtttgagcaagaaggaaaatcaggggcttctgttgggcaggtatgagattgggaaa<br>ttactaggacatggtacttttgctaaggtgtaccatgctagaaatgtgaagacgaatgagagcgt<br>agcgattaaggtgattgataaagagaagatcttgaaagtgggtcttattgatcacataaaacgtg<br>agatctcaattctcaagagagttaggcatccaaatatagttgaattatacgaagttatggctacca<br>aagccaaaatcttctttgtcatggagtatgtcaaaggcggtgagctcttcaataaggttgccaaag<br>gcaggcttaaggaagaagttgctcggaaatatttccagcagttaatttctgctgtcgctttttgtcatg<br>cccggggtgtttaccatagggacctaaagccagaaaatatactgttagatgaagatgggaatgt<br>caaagtttctgattttggtcttagtgccatttcagagcagataaagcaagatgggctcttccatacttt<br>ctgtggtaccccggcttatgtggcgccggaggtgttaggaagaaagggttatgatgcagccaaa<br>gttgacatttggtcatgtggggtgatactctttgtgctaatggcagggtatttgccatttcatgatcag<br>aatattatggctatgtacaagaagatttataggggtgaatttagatgtcccaggtggttttcacctga<br>attaactcggtttctaaagcgccttcttgatattaatcccgaaaccaggattactgttcaggagatta<br>tgaataataggtggttcaagaagggttttaaacatgttaaattttatattgaggatgataagttgtgta<br>gtataaatgatgatgagtatggtgggattgattattcgtctgatagatcagagtctgagtctgaaat |

| | | |
|---|---|---|
| | | agagatcagaaggaggtcagctagtttacctaggccagctagcttgaatgcatttgatattatttcattttcccgtggctttgatttgtccggtttgtttgaggagggaggagatggggcaaggtttgtatctggggctcctgtgcccaagatcataaataagttggaggaaattgctaaagttgtgagctttgcagtaaggaagaaggattgtaaagtgagtatggaggggtctaaggaaggtgcaaaggggccgttgacagttgcagctgagatatttgagttgacaccatcgttgagagttgttgaagtgaagaagaaaggaggagatagcttggagtatgaggagttttacaacagggaattgaagccaggattacagaatttagcacatgaagttggataccctgttaattcatcgtatttgccatcagatactgaataa |
| **SlCIPK13** | Solyc09g018280 | atgccagagatcatacttaataactatggaggcagtacatctttactatccactgccgcagcaatctacgctgaattaactccggctgagttacctccgccggaagacgatgttaattcaaaccttttgacaaatatgaactgggacaactcctgggttgtggtgcttttggtaaagtatatcatgccagagacttcaggacggcacagagtgtggccattaaagtcgtcagcaaacagaaaatactcaaaggtgggctaacggggcacgttaagagagaaatctctatcatgcgtcagttgcgtcatccccatatcgtacggcaacatgagattcttgctactaagaagaaaatctacttcgtcctggaattcgctaaaggaggtgaactcttcgccaagctagctaaaggccggttcagcgaagatctcagccgtcgatatttccagcagttaatctccgcagttggatactgtcactctcgcggggtctatcaccgtgatttgaaaccggaaaacttattactggacgaaaattgggacctgaaagttactgatttcgggttaagcgctgtcagggatcagatccgacctgacgggttgcttcatacactttgtggtacccctgcttatgtagcaccggagattctggggaagaaaggttatgatggtgctaaggtggatatatggtcatgtggtattattctctttgttttcaatgctgggtatttacctttcaatgacacaaatttaatgacgatgtatcggaaaatttacaaaggtgaatttcgttgtccgaaatggacctcgccggagctgaagaggttgttgacccgattacttgatatcaacccggttactcgtatcaccattgaggagattaagaatgacccgtggtttcaaacagggtatcaagaggtgatatcggtaaatcatcctttcgagttcaagtcatggtcgggtccaggttccaacggagaattccttaacgcatttgatattatctcgtattcatctggttttaatttatccagtttagttaaaggcaatggtggattcattattaaggaacaattcgtatcgagggagacgccggagaagataatcgggaaaattgaggaggtggcggaggtggaagggatgacggtggcagagaggagtggagcctcagtgaaggtggaggggcagaatggtaattttgtggtaatggtggtagttaaccgattaactgaagaactggtaatagttgaaattgagaaaaaggagaatgatggtgaaatttggaagaagaaattcaaaccggagataagtagatttgtttatcaagggtga |
| **SlCIPK14** | Solyc06g082440 | atgccggagaagcctatattcggaaaatatgagctaggaaagctactcggttgtggtgcatttgccaaagtgtaccacgctagagaaatcagcaacggaaagggcgtcgcaatcaaaataattaacaagagtaatattttgaataaagatggtattctcaacaatagaatcgagcgcgaggtgtgcattatgcggcagctgcaacatccgtacattgttagactctatgaagtgcttgctacgaagacgaagatctatttcgttatggaatacgtgaaaggaggcgaattattcaaccagatctccagtaaaagccgattcactgaggatctcagccggaaatgctttcagcaattgatttccgcggttaattactgtcattcacgtggaatttaccatcgcgatttgaaacctgagaatgttctaattgacgaaaatggagatctgaaagtttccgatttcgggcttagtgctacaacggatcaaattcaatcattcgacgggcttctacatacggtttgcggatctccggcgtatgtagcaccggaggttttgacgattagaggatacgacggagctaaaacagatatctggtcatgtggaattatgctattcgtgatgatcgccggttacttcccgttctacgatcagaatctgatgttaatgtacaaaaaaatctacaaaggcgaattccggtgcccgaaatggatctctccggacggtaagcggattctatctcgtcttctggatgtaaatccggcgaccagaattacgatcgaagaaatcatcagagatccatggttcagaaagggattgaaatttatcaaattctcagaagaggaagaaaatagtaagataaattcgttaacaaattctctgaatgcgtttgatataatttcgttctcccagggattagacatttctggattgttcaaatcgaacaatccggtggatgatttggagaggatagtggtggaagagtcgccggagggtgtaattgagagaattgaagaggtagggaagaaggagaatttcaggatgaagaggaaaaaagattggggaatcgatatgaaagtacaaaacggtaaattaacaatgaatttgaatttgaacgtatatcgattgatcgaacgattaacagtcgtcgaaattcaaaaaattgatggtaatgatgatttgtataaagacgtatggaggaataaattgaagcccgtaattttgagccaacgcggaacagagcttcacatttccgacagctaa |
| **SlCIPK15** | Solyc05g052270 | atggagaaaaaaggaaatgtactgatggaaaagtacgatttggggagattattaggtcaaggcaactttgctaaggtttattacggaaggaatcttgaaacgggacagagtgtagctgttaaggtaattgacaaagagaaggttattaaggccggactgattgaacagacgaagagagagagatatctgttatggcactagttgaacatccaaatgttctacagctatacgaggtcatggcaactaaatctaagatttatttagtcattgaacatgccaaaggcggtgagcttttcaaaaagttgacaaaggggaggctcaaggaaaaattagctaggaagtactttcagcaattgattagcgctgttgaatgttgccacagtcgagatgtttatcaccgtgatctgaaaccagaaaatgtgcttctagatgaggatggaaatcttaaggtatcagactttggattaagcgccttggctgagtctaagaggcaggacggcttactccacacaacatgtggaaccccagcttacgttgcacctgaggtgattagtcgaaaaggatatgatggcgccaaagctgatatctggtcttgtggggtgatcttatttgttttgctagccggttatcttccattccaagactcaaatcttatggagatttataggaagataaagcaggctgagttcaaatgtcctaattggttccctccagaggtgcggagattactttctaaaatccttgatccaaaccctcgtacaaggatttccatcgcaaagataaaggaaagctcatggttcaagaaaggtttcgagtctagaaacacggtaaccaaagtagaagaaaaggaaaagtttagcctagacgccaacgctacacataacagtatctctccctcagtttcaaagctaga |

| | | |
|---|---|---|
| | | gttgctaaaaccgacaaatctaaatgcatttgatataatctctctttcaagtggatttgacttgtccgg<br>tttattcataacaaaggatcagaaggaagacttgcagttcatttccgcaaagcctacctcttctatc<br>atgtccaaactcgaggaagttgggaggaatcttaagctggaagtaatgaagaaagaagctggt<br>tttatgagattagagggatcgagtgaaggtagatacgagactttgtccatcgatgctgaaatatct<br>gaaattactccgtccttccacttagttgagctgaaaaagtcgtatggtgataaggtagagtaccaa<br>aaattgctgaaacaagttataagacctgctcttgaggaaattgtttgggcttggcaaggcgagca<br>gccagctaatcgttag |
| **SlCIPK16** | Solyc09g083100 | atggatgtaaaacaaccccaaccacctccatttaccgtcaaaatcagaagaaccaccagttcc<br>gatggtagtggatctggatctatcattcttggaaaataccaattggttcgtctattgggtcgaggtag<br>ctttgctaaagtgtaccttggccgttgtttagacgataacactgaagttgctgttaaagttatagata<br>aatccagtactgccattgatgcttctatggagccacggattatccgtgaagtctccgccatgcgcc<br>gccttaatcaccacccgaatatcctcgaactttttgaagttatggcgacgaaaactaagatctattt<br>cgttatggaactagctcacggcggtgagcttttcacaaagcttaatcgccgtggccggttttctgaa<br>tccaccgccagattttatttccatcagcttgtctctgctttacatttctgccaccaaaacggcgttgctc<br>atcgtgatatcaagccacaaaatctactcctagacaaagagggtcatctcaaaatctccgatttc<br>ggactctccgccttgccggagcagtttgagaacggtcttcttcatacagcttgtggtacaccggcg<br>tatactgctccggaggtggtttacagaagagggtacgatggtgctaaggcggatgcttggtcatg<br>tggggttattctctttgtgttcctcgccggaagcttaccgttcgatgatagcaatttgccgagcatggt<br>taaatctatacaccgacgtgaatatacgtttcccgattgggtttcaaaatcagcccggaggataat<br>taaccggctacttgatcctaatccgaaaacgagatacggaattgaagagctcatgaacactcca<br>tggtttaagaaatcatcatcgatgaaaccagaacaaagcacgaagcaatttggtgagggaattt<br>tggagaaggaaagcaaacaaatggagagtataaatgcatttgatttaatttcaatgtgttcaggg<br>ttggatttatcgtcgatatttgaagaagaattgaacaagaaggagatgagatttacgacaaatgt<br>agaagttaaggtgatagaagagaaggtgatgaatgttggtaaagatgcaggatacagagtag<br>agaaaagaaagaatggtggaattgggttggtgaaagggagaagtgttttgttagttgaaatcttg<br>gagttggcaaaagagttgttgttagtggagttcaaagttgttaatggaggatcagaattcgaggat<br>cgtcaatgggaagaattgaaagctggattgaaagaagtagctgtttcatggtag |
| **SlCIPK17** | Solyc12g098910 | ATGGTACTGATTCAGCATCAACAACAGCAGCAGGAAGAAGAAAT<br>AGTACGAAGAGAAAGAGGGAAAAAGGGAATGCGAGTAGGGAAA<br>TACGAGCTTGGAAGAACACTAGGAGAAGGTAATTTTGGGAAAGT<br>GAAGTTCGCTAAGCATACAGATTCAGCCAAATCTTTTGCTATTAA<br>GATTTTGGAGAAGAATCGGATCCAGGATCTTCGAATTACTGATCA<br>GATAAAGAGGGAAATCCGAACTTTAAAAGTTCTCAAGCATCCAAA<br>TGTGGTTAGATTATACGAGGTATTAGCGAGCAAAACCAAGATTTA<br>TATGGTGTTGGAATATGTAAATGGTGGTGAATTATTTGACAGAAT<br>AGCTACTAAAGGTAAACTTTCAGAAACACTAGGCAGAAAATTATT<br>TCAACAATTAATTGATGGTGTTAGTTATTGCCATGACAAAGGTGT<br>CTTCCATAGGGACCTCAAGCTAGAGAATGTTCTCATTGATGGAG<br>GCAGAAACATAAAGATAACAGATTTTGGACTAAGTGCATTGCCTC<br>AACATCTTAGGGATGATGGCTTGTTGCATACAACATGTGGTAGTC<br>CCAACTATGTTGCTCCTGAAGTTCTTTCTAACAGAGGCTATAATG<br>GTGCAACATCAGATACTTGGTCTTGTGGTGTCATTTTATATGTCA<br>TCCTCACTGGCTTTTTACCCTTTGATGATAGAAATCTTGCAGTGC<br>TTTATCAAAAGATTTTTAAAGGGGATGCTCCAATACCAAAGTGGT<br>TATCACAAGGAGCAAAGAATCTTATAAAGAGGATTCTTGATCCAA<br>ATCCACAAACTAGAATAACAATGGCAGAGATTAAAGAAGATGAAT<br>GGTTTAAACAAAACTATACTCCTACAAATCCTGATGAAGAAGAAG<br>TGGAAAGTGATGATACATCCTCAGATGATGAAGTGTTGACAATAC<br>ATGAAGCACCACTTGACATAGAAAGAAATCCAGAATCACCTTCTG<br>TCATCAATAATGCCTTTCAACTTATAGGAATGTCCTCATGCCTTGA<br>TCTATCTGGATTTTTTGAAAATGAGGATGTTTCTGAGAGGAAGAT<br>CAGGTTCACATCTAATCTCTCTCCAAAAGAACTGCTAGATAGGAT<br>TGAGAATCTAGCTGTTCAAATGGGATTTCAAGTCCAGAAAAAACC<br>TGGAAAGTTAAAAGTATTGCTAGAGAACAAAGGTCAAAAAACACA<br>AGCAAGTCTTTCAATAGTAGTAGAGGTTTTCGAGATTAGTACATC<br>CTTGTATGTTGTAGAGTTACAGAAATCTTCCGGGGATTCTGCAGT<br>ATATAGACAGTTGTGTAATAAATTATCAGATGAACTGGGAGTCCA<br>GCAAAGTGAAGAACTACTGACTAATCTAATTGTGAAAGATGGAAA<br>TCATACCAGTTAG |
| **SlCIPK18** | Solyc09g042660 | atggagagtaagaggagtaaggggagtagtatactaatggaaaggtatgagttagggagattg<br>ttaggacaaggcacctatgctaaggtccatcatgcgagagacgtcaagactagcatgaatgtg<br>gctattaagattattgataaggagaagattgtaaaagttgggatgattgatcaaatcaaatgcga |

| | | |
|---|---|---|
| | | gatttcagcgatgaaattggttaggtattctaatattgtgcagctctatgaggtgatggccagcaaa<br>aagaagatttattgtgtaatggaatatgtcaaaggaggcgagttgtataacaaagtgagtaaag<br>ggaagttaaaggaagatgttgcaaggaagttatttcagcaattgattagtgctattgatttctgtcat<br>agtagaggtgtttatcaccgagacctaaaaccggagaatctattgcttgatgaaaatgggaatct<br>aaagatttcagattttggattgagtgcccttgcagattcgaaatgccaagacggtttacttcatacta<br>aatgtgggacaccagcttatgttgcaccagaagtgataagcaagagaggctatgatggtgcga<br>aagctgatatttggtcatgtggggtgatattatatgtccttttggctggatatctccctttccaggattca<br>aaattaacggagatgtacagaaagatcggtaaggctgagttcaaatgtcctaactggttccctcc<br>tgatgctcgcaggctgatatcaaaaatattaaatccaaatcctagcacaagaatttccatttctaa<br>gataatggagaattcttggttcaagaaagggttgcagctgaaacctataatcgctgatgctgactc<br>aaagcctgaagaagcagcaattgatgatgcagtttttggtatcggtgagagtaccgactcaata<br>acagaaccaaagccagaactggtcaagcctgcaTcattgaatgcatttgatattatatctttctca<br>acaggttttgacttgtctgcctttttgaagaaagagataaaaagagggaagttatatttacatcaa<br>agcaaccggcgaaaacaatcatctcaaagctagaggatgtagctaggcgtttaaagttgaagg<br>tgatgaaaaaggatggagggttattgaagttggaaggatctaaagaaggaagaaaaggggtg<br>ttgtctattgatgcagaaatctttgaggtaactccaaattttcactttgttgagatgaagaaatcgaat<br>ggagatacgatagagtacaagaagacgatgatgacggatgtaagaccagcattggaagata<br>ctgtatggacttggcaaggtgacgaaccgcgtcctcagcagctagcagaggaagaaattttgc<br>agacttatcaggggtactcatttgacaataattcacctcagaagaatgcataa |
| **SlCIPK19** | Solyc03g006110 | ATGATGAGCATGAAGAGCGCGAAAACGGAGAAAAAAGGGACAAT<br>TTTGTTGCATAAATATGAGATAGGTAAATTGTTAGGGCAAGGTAC<br>ATTTGCCAAAGTTTATTATGCAAGGAACCTAAAAACAGGACAAAT<br>TGTAGCTGTTAAGGTAATTGACAAAGAAAAAGTGATGAAAGTTGG<br>CTTAATTGACCAAATCAAACGTGAAATTTCTGTTATGAGATTAATT<br>AGGCATCCAAATGTTGTCGAACTTTATGAAGTAATGGCTAGCAAA<br>ACAAAAATTTATTTTGCAATGGAATATGTTAGAGGTGGTGAATTAT<br>TCAATAAGGTTGCTAAAGGGAGGCTTAGGGAATCAGCAGCGCGA<br>AAATATTTTCAACAATTAATCGCGTCTGTTGATTTCTGTCATAGTC<br>GAGGGGTATATCATAGGGACCTGAAACCTGAAAATATACTCCTG<br>GATGAAACTGGAAACCTGAAAGTTTCTGATTTTGGGTTGAGTGCT<br>CTTTTCGATACGAAAAGACAAGATGGACTCCTCCACACGACGTG<br>TGGAACTCCAGCATATGTCGCGCCAGAGGTCATAAATAAGAGAG<br>GATATGATGGCGAAAAGGCGGATATTTGGTCATGTGGGGTGATT<br>TTGTTTGTGTTGTTAGCTGGTTATCTTCCATTTCATGACACAAATT<br>TACTGGAAATGTATAAGAAGATTACTAAAGGGATATTTAAATGCC<br>CTGAATATTTTCCATATGAAGTGAAAAAACTACTCTTAAGAATTCT<br>TGATCCAAATCCAATTTCAAGAATTACATTAGCTAAGCTAATGGA<br>CAATAATTGGTTCAAGAAAGGATTCAAACAAATTGATAAACCATTC<br>ATTTTGGATCAAGATCACGACGACGATTCACCTCGTAGTGTTTTT<br>GATATGGTGGATGATTCGGATGCAGAGTGTTCTTCACGACACAA<br>AGAAGAGAGCTCATCAACAATAATGAAGCCTACTTGTTTGAATGC<br>TTTCGATATCATTTCACTTTCTCCTGGTTTTGATCTCTCTAGCTTG<br>TTCGAGAAAGATAAGAGTCATAGGTCGGATGCTAGATTCACGAC<br>ACAAAAGTCTGCATCCACTATCGTGTCAAGGCTAGAAGAAGTAG<br>CTTCAATGGGGAGCTTTAAGGTGAAGAAAAAAGACGGGACGGTG<br>AAAATGCAAGGAAGCAAAGAAGGGAGAAAAGGGCAATTAGCAAT<br>TGATGCAGAGATATTTGAGATAACTCCAGCTTTCCATGTTGTGCA<br>AGTGACAAAAAAATCAGGTGACACTGCAGAATACAGGAACTTTTG<br>TGATCAAGGATTAAAGCCATCATTAAAAGACATAGTTTGGACATG<br>GCAGGGCAATGAGCAATTACAACAAGTGGAAAATCAAGAAAACA<br>AGACTTAA |
| **SlCIPK20** | Solyc02g072540 | atgacttcgaagaacgataagaaaggcaatattttgatgcaaaggtatgagattgggaaattgct<br>tgggcagggtacatttgcaaaagtataccatgctagaaatctcaaaacagggcaaagtgttgct<br>attaaggtgattgacaaagagaagattatgaaggttgggttaattgatcaaacgaaacgtgaaa<br>tctccgtcatgaggctaatcaaacacccaaatattgtccagctctatgaggttatggcgagcaaa<br>acaaaaatatattttgctatggaatatgttagaggtggtgaacttttcaataaggttgctaaaggca<br>gacttaaagaagatgctgcaagaaaatactttcaacagttaatcgctgcagtcgatttctgtcata<br>gccgtgatgtctaccaccgtgatctcaagcctgaaaatctcctccttgacgaaggtggcaatctg<br>aaagtctcggattttggtctgagtgcattgtttgattcgaaaaggcaagatggtctactccacacga<br>cgtgtggaacaccggcttatgttgcaccagaggtgatcaataagagaggttatgatggtgaaaa<br>ggctgatatttggtcatgtggtgttgttttgtttgtcctgttagcgggttatttgccattccatgaccaaaa<br>tcttatggaaatgtacaaaaaaaataagcaaagcagaattcaaatgtccacaatggttccatcca |

| | | |
|---|---|---|
| | | gaggtaagaaagttactctcaaggattcttgatccaaatccaggatcaagaatcaccttaattaa<br>gctcatggagaattattggttcaaaaaaggattcaaacaagttgataaaaccccaaattcaggg<br>aaagatcaacgtgaatcccctcgtagtgttttcgatattgaggacaataattcagacggagaggg<br>accttccaatcgccaaaagaatcaagattccacaacaatgaagcctacttgcttaaatgcatttg<br>atataatctctctttcccctggtttcaatctttctggattattcgaaaaggagaaagagagaagatca<br>gaagcgcgattcacagcgaaaaaaccagcttctataatagtgtcaaaattggaagaagtagctt<br>caaatgagagttttaatataatgaaaaaagatggtacagtgacaatgcagagtaacaaagaa<br>ggaaggaaagggcaactagcaattgatgcagagatattcgaaattacgccttcttttcatgttgtg<br>gaagtgagtaaaaaatcaggggatacaatggaatacaagaaattctttgatcagggattgaaa<br>acatcacttaaagatattgtttggacatggcaagatggtgaacaacaacaacaagttgaaaatc<br>aagaaagaatttga |
| **SlCIPK21** | Solyc12g010130 | atggggacagaagaaaaatgtgctgttttgtatggtaaatatgagctcggacgagttttgggtcaa<br>ggttctttcgctaaagtataccatgctcgtaacgtggtaacaggggagaatatagcaatgaaagt<br>tgttggaaaagagaaggtgataaaagtaggaatgatggagcaaatcaaaagagaaatctcc<br>gtcatgaaaatggtgaaacatccaaacatagtcgagcttcacgaagttatggcaagcaaaacg<br>aagatttacttcgccatggaatacgttaaaggcggtgaattgttcgaaaaagtagctaaaggtaa<br>gcttagagaagacaatgctcgagggtatttccagcagttgatttctgcaattgatttttgtcatagcc<br>gtggtgtttatcacagagatttaaagcctgagaatttgctgttagatgaagaaggcaatcttaaggt<br>aaccgatttcggacttagtgcttttactgatcatttaaggcaagatggtttattgcatacaacttgtggt<br>actcctgcttatgttgcccctgaagtacttggtaatcatggatatgatggtgcaacatcagatatttg<br>gtcatgtggagtgattctttatgtgcttttagctggtttttaccatttcaagatgataatatcatggctat<br>gtataagaaaattcacaagggtgatttcaagtgtccaccttggatgtcatctgatgcaaagaagtt<br>gattgtaaagatgttggatccgaatccgagaactcgaattactgcttcaaagattatggagtctaa<br>ttggttcaagaagactgtgccaaggactttgaggagtaaggtggaggaagagttttctcatgtgg<br>gagacgaagattgtgtagggaaggcgaaaaagattgagtctttgaatgcttttcatatcatttcatt<br>gtcggaggggtttgatttgtctcctttgtttgaggaaaagaagaagaaggagaaggaacagttg<br>agatttgcaacaacaaagccagctagtagtgtgatttcgaagctcgaggaagtagctaaaacgt<br>cgaaattcagcttgaaaaggagtgattctagtgttagattgcaggggcaagagagtgggagga<br>aagggaagttgggaatatctgctgatatatttgctgtgacaccttcatttctggttgtggaggtgaag<br>aaagcttgtggtgatacattggaatataatcaattctgcagcaaagagcttaggccagcactcaa<br>ggacattgtttggaaatccgcacctgagaatccaacaattgcttga |
| **SlCIPK22** | Solyc06g007440 | atggctttattctcatctgaaaccacgatattcaacaaatacgagctaggaaaactcttaggatgt<br>ggcgcatttgccaaagtgtaccacgctagggacattagagatggccgtagcgtagcgattaaa<br>attattaacaaaacaagagttagcaacgcgattttaatggcgaatatcaaacgcgagatctcaat<br>tatgagacttttacgacacccacacattgtcaaacttgacgaagtgttggccaccaagacgaag<br>atctacttcgtcatggaatttgttaaaggaggcgaattattcgccaaaattgctaaagctggaaaa<br>tttcccgaagatcaaagtcggaaaattttccagcaattaatttctgctgttcgttattgtcattctcggg<br>gtgtgtatcatcgtgacctgaaacctgaaaatttacttatcgatgaaaatggtgatttgaaagtttct<br>gattttgggctaagtgcgttaacggaacaagtacaacaagatgggttgttacacacgctttgtgg<br>gaccccatcttacgtggcaccagatgttttaacaaaaaaaggatacgatggagctaaggcgga<br>tatttggacatgtggaatcatcttgtttgtgttaaatgctggatatttgccatttcatgattcaaatcttat<br>gggaatgtaccataagattatcaatggagaatacaagtgtccaaaatggatgtcatctgagttaa<br>agcgatttttacgtagacttcttgatactaatccaatgacaagaataaatgtagaagaaatcatga<br>atgatgcatggtttaagaaaggcttgaaacgcgttaaattctgtgaggaagatggtaagcctcag<br>agacgaattcagaattcagattcaacagatcaggatgagcctagcgcggaaatagaagaggt<br>caaggatggagaagacaagaagaagaattcttatttgaatgcttttgatataatttcattctctatgg<br>gattagacctctccggattgttaaatgacggttttaaccctttagaggatttcgaaagattagttgttg<br>aagaatcattggagatagtaatggagaaagtagaagaaatggcaaagaaggagaatatgag<br>gttaaagaagaaaaaaagaagggggaatagagttgaagaaaggtaaattgattatgaacatgg<br>aaataagtagagtgatagatggatttgttgtcgttgaggttcgtcgaaaagtcggtgatactgatac<br>atacaaagaaatattgaagaataatataaagccaataattctcggccaagagccgatggaagt<br>attggttccgacagagatggatccataa |
| **SlCIPK24/<br>SlSOS2** | Solyc12g009570 | atgaagaaagtgaagagaaagcttgggaagtatgaagttggcagaactattggtgaagggac<br>atttgccaaggttaagtttgcacgaaacaccgagactggagagaatgttgccattaaagtcttgg<br>ccaaaagtaccattcttaagcatagaatggttgaacagatcaaaagagagatatctataatgaa<br>gattgtcagacatccttgcatagttcgacttcatgaggttttagctagccagacaaaaatatatatc<br>gttcaggagtttgtcactggaggagagctttttgataaaattgttcatctaggtaggctttctgaggat<br>gaagcaaggagatacttccaagaactcatagatgcaattgctcactgtcacagcaagggtgttt<br>accacagagatttgaagcctgaaaatttgcttcttgatttccaagggaacttaaaaatttctgacttt<br>gggcttagtgcattgcctcaacaaggagtcgagctcctctataccacttgtgggactccaaattat<br>gttgcacctgaggtgttaggtaaccgaggttatgatggtgctgctgcggatgtgtggtcatgtggta<br>tcatcctttatgttctgatggcaggatatcttccatttgatgagacagaccttcctaccttgtatacaaa |

| | | |
|---|---|---|
| | | gatcaaggctgctgaattttcctgtccattttggttttctcctggtgcaacatcgttgattcaaaaaatc<br>attgatccaaatcctcaaactaggatcaagattgatggaataaaacgagacccttggttccgga<br>aaaactacagagctgttaaagctaaagcagatgaagtagtgaatcttgatgatatccatgctgtg<br>ttcgacgacattgaggatgcatttgtcagtgaaaaatcagaagatagtgagagtggtcccttggt<br>aatgaatgcatttgagatgataacactatctcagggattaaatctatccgctttgtttgacagacgtc<br>aggattatgtcaagcgtcaaactcgatttatttcccgccaacctgctaaagtcgtcattgaaactatt<br>gaagcagctgcagagtcgttgggtcttaaggtccacacacgcgattacaagacaagaattgag<br>ggggtaacggcaaatagggctggtcaatttgctgttgtgctggaggttttccaagtagccccttcc<br>ctttttatggttgatgttagaaaggctgctggggacactcttgaatatcacaagttctacaaaacctt<br>ctgcacgaaaattgacgacgtcatttggaaaccgaaggaaggcatgtcaaatgctgttctgctta<br>ggacaaggactcgctga |
| **SlCIPK25** | Solyc06g068450 | atggaagatcaaattgatcaatcgaagcaagaaatggatcatttaatggatggttcgagaagtat<br>tatattcgggaagtacgagatggggaggctattaggccaaggaacatttgcaaaggtttattata<br>gcagagatatcaaaacttgtgaaagcgtagcgattaaagtaatcaacaaagatcatgttaaaa<br>gagaaggaatgatggagcaaatcattcgagaaatttcaatcatgcgattagttcgacatccaaa<br>catcgtggaaatcaaagaagttatggctacgaaacaaaaaatctttgtagttatggaatatgttaa<br>agggggtgagcttttcgcaaaagttgctaatggcaaacttaaggaagatgttgcaagaaaatac<br>tttcagcaattgattagcgctgttgatttctgccatagtcgcggtgtatttcaccgcgatttgaaacct<br>gagaacttactccttgatgaaaatgaaaacttgaaggtgtctgattttgggttatcagctttgtctga<br>gcaattgaggagcgatggtttgcttcatacacggtgtggaactccagcttatgttgcccctgaagtt<br>ttgaggaaaaaagggtatgatggtgctaaatctgatatttggtcttgtggggttattttatatgttctttt<br>agctggatttctaccatttaaacacgagaatttgatgaagatgtatcgaaaagtctttaaaggtga<br>atatgaatttcctccatggttttcccctgaagctaagaaactgatatcaaagcttctagtagctgatc<br>cagaaaaaagaattacaatttcagctgtaacaaaagtcccttggttatcaaagaattcaatcgat<br>cttcttctttttcttccatcgaagaaaacaacacagatcagaaacaggaaaccaaagaaccaag<br>tttgggagctagatcaaaatcagccccaccttttttacaacgcatttgaattcatatcgtcaatgtcttc<br>aggtttcgatttatcgagccttttttgaaggtaaaaagaaatccggttctcttttcacatcgaaatgctc<br>tgcttcaacgatcatgtcaaaactcgaatctttagcgaagaaggtgaattttcagattgtatccgct<br>aaggaattcaaggtgaagatgcaaggaacatcgaatgggcgaaaagggaaattgtcagtcat<br>ggctgaagttttcgaagttgcaccacaagtggcgattgttgaattctcaaaatctgccggagaca<br>ccttagaatataaaaagttctgtgaagaagatgtacggcctcccccttaaagatattgtttggacatg<br>gcaaggtgagaataatggccgtgattaa |
| **SlCIPK26** | Solyc11g062410 | atgaatcaggcaaaaatcaagcgcagagttggtaaatatgaaatgggaaggacaattggtga<br>gggaacatttgctaaagttaagtttgcgaggaattcggagacaggagaagctgtagcgatcaa<br>gattcttgataaggataaggtccttaagcacaaaatggctgagcagataaagcgggagatagc<br>tacaatgaagttaattagacatccacatgtcgttcagttatacgaggtgttggctagcaagacaaa<br>gatattcattgttctggagttcgttacaggcggagagctctttgacaaaattgtaaatcatggacgg<br>atgcatgaaaaagaagcaaggaaatactttcagcagcttattaatgctgttgattattgtcatagc<br>aggggagtctaccatagagatttaaagcctgagaatttactcttggatgtcaatggaaacctcaa<br>ggtttctgattttggattgagtgctctgtcccagcaagtcagggatgatggcttactccacaccacat<br>gtggaactcccaactatgttgctcctgaggtactcaacgatcatggatatgatggtacaaccgcg<br>gatctgtggtcgtgcggagtcatactctttgtattgcttgcaggttacttgccttttgatgactccaatct<br>tattaacctgtataaaaaaaatctctgctgctgaatttacatgtcctccttggatgtcttttggtgctatga<br>agttaattactcgcatcttggatccgaaccctatgacacgtatcaccatctcagaaattctggagg<br>atgagtggttcaagaaagattataaatctcccatttttaatgagaaaggagatgccaacctggat<br>gatgttgaagctgtcttcaaggattctgaagaacatcatgtaacagagaaaaaggaagagcag<br>ccaactcccatgaatgcattcgagttgatttcaatgtcaaaaggactcaaccttgggaatctcttcg<br>atgaacaggaatttaagagagaaacaaggttcacatctaaatgctcggccaatgaaattatca<br>gtaagattgaagaagctgcaaagccccctcggttttgatgttcacaaaaagaactacaagatgag<br>acttgaaaatgtcaaagctggaagaaaagggaaccttaatgttgccactgaggtatttcaagttg<br>ccccttctcttcatatggttgaagtccgaaaggcaaaaggagatactttggaattccacaagttct<br>acaagaatctttcgactagtctagaggatgtagtgtggaaaactgaagaggacatgcaagcta<br>ggtag |
| **SlCIPK27** | Solyc06g050270 | ATGGTGAAAAATGGAAATGTAGTGATGCAAAAATATGAATTGGGG<br>AGATTATTAGGTCAAGGCAACTTTGGTAAGGTTTATTATGGAAGA<br>GATCTTGAAAGCGGACACAATGTAGCCATTAAAGTAATTGATAAA<br>GAGAAGGTTCAGAAAGCTGAATTGACTGAACAGACGAAACGAGA<br>GATATCAGTTATGGCAATGGTCAAACATCCACATGTTGTGCAGCT<br>ATACGAGGTCATGGCAACTAAGAGTAAGATTTACTTTGTGATCGA<br>ACAGGCCAAAGGTGGCGAGCTTTTCAACAAACTGACAAAGGGCA<br>GACTCAAGGAAGACGCTGCTAGAAAGTTGTTTCAGCAACTGATC<br>AATGCAGTTGAATTTTGCCACAGCCGAGGTGTTTATCACCGTGAT |

| | | |
|---|---|---|
| | | CTCAAACCAGAAAATCTCCTACTGGATGAGAATGGAAACCTAAAG<br>GTCTCAGACTTTGGTTTGAGTGCATTAGCTGAGTCTAAGCGACAA<br>GACGGGTTACTTTACACAACCTGTGGTACACCAGCATATGTTGCT<br>CCTGAGGTGATCGGTAGAAAAGGATATGAGGGTGCCAAAGCTGA<br>CATCTGGTCTTGTGGAGTGATTTTATTTGTCTTGTTGGCTGGTTA<br>TCTTCCATTCTATGACTCAAATCTTATTTATCTGTATAGGAAGATA<br>TGCAAGGCGGAGTACAAATGTCCTAATTGGTTCCCTCTAGAAGT<br>GCGTAAACTTCTCTCTAGGATCTTCGACCCAAACCCTCATAAAAG<br>GATTTCAATAGCCGAAATAAAGGCAAGCTCCTGGTTTAAGAAAGG<br>ACTGGGATCTAAACAAGTAGTGAACCAAAATGTTATTGCAGATGG<br>TGATGCTGTTTCCAGTTTGGATAATACCAAGTTAGAGTTGATAAA<br>ACCTGCAAGCGTTAGTGCATTTGATATCATCTGCTGGTTTAATTT<br>GTCTGGTTTATTCATAGACGATGATCAAAAGGAGGAGCTGCGATT<br>CACATCAGTGAAACCTGTCCCAGTCATCATATCTAAGCTTGTGGA<br>AGTTGGCAAGAGTCTGAACCTTGAAGTAAAGAAGAAAGAAGTTG<br>GATTTCTTATGTTGGAGGGATTAAATGAGAGCAGATATGAAACCG<br>TGTGCATTGGCGTGCAAATCTTTGAAATTTCTGTATCCCGTTACT<br>TCATTGAGCTGAGTAGGTCAAGTGGTGATGCGATTGATTACCAAA<br>ACATGTTGACGCAAACTATCAGACCAGCTCTTGATGAAATTGTTC<br>AGGCTTGGCAAGGTGTGCAATCTCATCAACAATCACTAAAATGA |
| **SlCIPK28** | Solyc06g050280 | ATGGTGAAAAAGGGAAATGTAGTGATGCAAAAATATGAATTTGGG<br>AGATTATTAGGTCAAGGCAACTTTGGTAAGGTTTATTATGGAAGG<br>GATCTGGAAAGAGGACAGGCTGTAGCAATTAAAGTAATTGATAAA<br>GAGAAGGTTCAGAAATCTGAATTGACTGAACAGACGAAACGAGA<br>GATATCCATTATGGCAATGGTCAAACATCCACATGTTGTGCAGCT<br>ATACGAGGTCATGGCAACTAAGAGTAAGATTTACTTTGTGATCGA<br>ACAGGCCATGGGTGGCGAGCTTTTCAACAAACTGACAAAGGGCA<br>GACTCAAGGAAGACGCTGCTAGAAAGTTGTTTCAGCAACTGATC<br>AATGCAGTTGAATTTTGCCACAGCCGAGGTGTTTATCACCGTGAT<br>CTCAAACCAGAAAATCTCCTACTTGATGAGAATGGAAACCTAAAG<br>GTCTTAGACTTTGGTTTGAGTGCATTAGCTGAGTCTAAGCGACAA<br>GACGGGTTACTTTACACAACCTGTGGTACACCAGCATATGTTGCT<br>CCTGAGGTGATCTGTAGAAAAGGATATGACGGTGTCAAAGCTGA<br>CATCTGGTCTTGTGGAGTGATTTTATTTGTATTGTTGGCTGGTTAT<br>CTTCCATTCTATGACTCAAATCTTATTTATCTGTATAGGAAGATAT<br>GCAAGGCGGAGTACAAATGTCCTAATTGGTTCCCTTTAGAAGTG<br>CGTAAACTTCTCTCTAGGATCTTCGACCCAAATCCTCATAAAAGG<br>ATTTCAATAGCCAAAATAAAGGCAAGCTCCTGGTTTAAGAAAGGA<br>TTGGGATCTAAGCAAGTAGTGAACCAAAATGTTATTGCAGATGGT<br>GATGCTGTTTCCAGTTCAAATAATACCAAGTTAGAGTTGATAAAA<br>CCTGCAAGCATTAGTGCATTTGATATCATCTGCTGGTTTAATTTGT<br>CTGGTTTATTCATAAACGATGATCAAAAGGAGGAGCTGCGATTCA<br>CATCAGTGAAACCTGTCCCAGTCATCATATCCAAGCTTGTGGAAG<br>TTGGCAAGAGTCTGAACCTTGAAGTAAAGAAGAAAGAAGTTGGA<br>TTTCTTATGTTGGAGGGATTTAATGAGAGCAGATATGAAACCGTG<br>TGCATTGGCGTGCAAATCTTTGAAATTTCTGAATCCCGTTACTTC<br>ATTGAGCTGAGTAGGTCAAGTGGTGATGCGATTGATTACCAAAA<br>CATGTTGACGTTAACTATCAGACCAGCTCTTGAGGAAATTGTTCA<br>GGCTTGGCAAGGTCTGCAATCTCATCAACAATCACTAAAATGA |
| **SlCIPK29** | Solyc06g050290 | ATGGCAATGGTCAAACATCCACATGTTGTGCAGCTATACGAGGT<br>CATGGCAACTAAGAGTAAGATTTACTTTGTGATCGAACAGGCCAT<br>GGGTGGCGAGCTTTTCAACAAACTGACAAAGGGCAGACTCAAGG<br>AAGACGCTGCTAGAAAGTTGTTTCAGCAACTGATCAATGCAGTTG<br>AATTTTGCCACAGCCGAGGTGTTTATCACCGTGATCTCAAACCAG<br>AAAATCTCCTACTTGATGAGAATGGAAACCTAAAGGTCTCAGACT<br>TTGGTTTGAGTGCATTAGCCGAGTCTAAGCGACAAGACGGGTTA<br>CTTTACACAACCTGTGGTACACCAGCATATGTTGCTCCTGAGGTG<br>ATCGGTAGAAAAGGATATGAGGGTGCCAAAGCTGACATCTGGTC<br>TTGTGGGGTGATTTTATTTGTCTTGTTGGCTGGTTATCTTCCATTC<br>TATGACTCAAACCTTATTTATCTGTATAGGAAGATTTGCAAGGCG<br>GAGTACAAATGTCCTAATTGGTTCCCTCTAGAAGTGTGTAAACTT<br>CTCTCTAGGATCTTCGACCCAAATCCTCATAAAAGGATTTCAATA |

| | | |
|---|---|---|
| | | GCCAAAATAAAGGCAAGCTCCTGGTTTAAGAAAGGATTGGGATC<br>TAAGCAAGTAGTGAACCAAAATGCTATTGCAGATGGTGATGCTGT<br>TTCCAGTTCAAATAATACCAAGTTAGAGTTGATAAAACCTGCAAG<br>CGTTAGTGCATTTGATATCATCTGCTGGTTTAATTTGTCTGGTTTA<br>TTCATAAACGATGATCAAAAGGAGGAGCTGCGATTCACATCAGT<br>GAAACCTGTCCCAGTCATCATATCCAAGCTTGTGGAAGTTGGCA<br>AGAGTCTGAACCTTGAAGTAATGAAGAAAGAAGTTGGATTTCTTA<br>TGTTGGAGGGATTAAATGAGAGCAGATATGAAACCGTGTGCATT<br>GGCGTGCAAATCTTTGAAATTTCTGTATCCCGTTACTTCATTGAG<br>CTGAGTAGGTCAAGTGGTGATGCGACTGATTACCAAAACATGTT<br>GACGCAAACTATCAGACCAGCTCTTGAGGAAATTGTTCAGGCTT<br>GGCAAGGTCTGCAATCTCATCAACAATTACTAAAATGA |

**Table S2.** Primers used in this study.

| Primer name | Sequence | Use |
|---|---|---|
| qSlEF1aF | GGCGGTGGCGAGCAT | qPCR |
| qSlEF1aR | AAACCAAGGCACCTCAACAAA | qPCR |
| SlCIPK15_qPCR_F | CACAACATGTGGAACCCCAG | qPCR |
| SlCIPK15_qPCR_Rv | AGATCACCCCACAAGACCAG | qPCR |
| SlCIPK26_qPCR_F | AAGCACAAAATGGCTGAGCA | qPCR |
| SlCIPK26_qPCR_Rv | TTGCTAGCCAACACCTCGTA | qPCR |
| SalI F SlCIPK26 | CCCGTCGACATGAATCAGGCAAAAATCAAGC | Cloning BIFC |
| XmaI R SlCIPK26 | ACTCCCGGGCTACCTAGCTTGCATGTCCTC | Cloning BIFC |
| SalI F SlCIPK15 | CCCGTCGACATGGAGAAAAAAGGAAATG | Cloning BIFC |
| XmaI R SlCIPK15 | ACTCCCGGGCTAACGATTAGCTGGCTGCT | Cloning BIFC |
| F sgRNA 1 SlCIPK26 | GTGCAGCAAAAATCAAGCGCAGAGT | Cloning sgRNAs CRISPR |
| R sgRNA 1 SlCIPK26 | AAACACTCTGCGCTTGATTTTTGCT | Cloning sgRNAs CRISPR |
| F sgRNA 2 SlCIPK26 | GTGCATCCGCACGACCACAGATCCG | Cloning sgRNAs CRISPR |
| R sgRNA 2 SlCIPK26 | AAACCGGATCTGTGGTCGTGCGGAT | Cloning sgRNAs CRISPR |
| F sgRNA 1 SlCIPK15 | GTGCACTTCGTCTGTTCAATCAGTC | Cloning sgRNAs CRISPR |
| R sgRNA 1 SlCIPK15 | AAACGACTGATTGAACAGACGAAGT | Cloning sgRNAs CRISPR |
| F sgRNA 2 SlCIPK15 | GTGCAGCGCCTTGGCTGAGTCTAAG | Cloning sgRNAs CRISPR |
| R sgRNA 2 SlCIPK15 | AAACCTTAGACTCAGCCAAGGCGCT | Cloning sgRNAs CRISPR |
| gSlCIPK15 p1 F | GGGTGAGGTCTGCAACAAGG | Genotyping of mutant plants |
| gSlCIPK15 p1 rv | GTAATCTCCGCACCTCTGGAG | genotyping of mutant plants |
| gSlCIPK26 p1 F | GGAAGCAACCTCTCGATCTCC | genotyping of mutant plants |
| gSlCIPK26 p1 rv | GTAGCTATCTCCCGCTTTATCTG | genotyping of mutant plants |
| gSlCIPK26 p2 F | CGTGGAGCTTGGGACAATGTC | genotyping of mutant plants |
| gSlCIPK26 p2 rv | GCTGCATAGGAGACACGTGC | genotyping of mutant plants |
